# Additive effect of contrast and velocity proves the role of strong excitatory drive in suppression of visual gamma response

**DOI:** 10.1101/497214

**Authors:** E.V. Orekhova, A.O. Prokofyev, A.Yu. Nikolaeva, J.F. Schneiderman, T.A. Stroganova

## Abstract

Visual gamma oscillations are generated through interactions of excitatory and inhibitory neurons and are strongly modulated by sensory input. A moderate increase in excitatory drive to the visual cortex via increasing contrast or motion velocity of drifting gratings results in strengthening of the gamma response (GR). However, increasing the velocity beyond some ‘transition point’ leads to the suppression of the GR. There are two theoretical models that can explain such suppression. The ‘*excitatory drive*’ model infers that, at high drifting rates, GR suppression is caused by excessive excitation of inhibitory neurons. Since contrast and velocity have an additive effect on excitatory drive, this model predicts that the GR ‘transition point’ for low-contrast gratings would be reached at a higher velocity, as compared to high-contrast gratings. The alternative ‘*velocity tuning*’ model implies that the GR is maximal when the drifting rate of the grating corresponds to the preferable velocity of the motion-sensitive V1 neurons. This model predicts that lowering contrast either will not affect the transition point or will shift it to a lower drifting rate. We tested these models with magnetoencephalography-based recordings of the GR during presentation of low (50%) and high (100%) contrast gratings drifting at four velocities. We found that lowering contrast led to a highly reliable shift of the GR suppression transition point to higher velocities, thus supporting the excitatory drive model. No effects of contrast or velocity were found for the alpha-beta response power. The results have important implications for the understanding of the neural mechanisms underlying gamma oscillations and the development of gamma-based biomarkers of brain disorders.

## Introduction

Gamma-band oscillations arise from a precise interplay between excitation (E) and inhibition (I) (Buzsaki and Wang, 2012; Vinck et al., 2013) and their parameters are affected by external and internal factors that can shift the E-I balance (Chen et al., 2017; Hadjipapas et al., 2015; Jia et al., 2013; Salelkar et al., 2018; Sumner et al., 2018). These oscillations are most prominent in the primary visual cortex (V1) and can be induced in both humans and animals by a range of visual stimuli. Human visual gamma oscillations recorded with Magnetoencephalography (MEG) have high within-subject reproducibility (Hoogenboom et al., 2006; Muthukumaraswamy et al., 2010; Tan et al., 2016) and are strongly genetically determined (van Pelt et al., 2012). This makes the properties of gamma oscillations potentially useful signatures of the constitutionally up- and down-regulated E-I balance in the human visual cortex. Various attempts have been made to relate the individual values of power and frequency of the visual induced gamma response (GR) to indirect measures of the excitatory state of visual circuitry, such as concentration of GABA and glutamate neurotransmitters (Cousijn et al., 2014; Edden et al., 2009), GABAa receptor density (Kujala et al., 2015), or performance in a visual orientation discrimination task (Edden et al., 2009), but the link between gamma activity and the excitatory and inhibitory processes in the visual cortex is still far from clear.

The relationship between the excitatory state of the visual cortex and the properties of gamma oscillations can be clarified through manipulations of the intensity of excitatory input. Interestingly, the intensity of visual stimulation has a different impact on the frequency and power of induced gamma oscillations in both monkeys and humans. The frequency of gamma oscillations nearly linearly increases with increasing visual input strength (Hadjipapas et al., 2015; Jia et al., 2013; Murty et al., 2018; Orekhova et al., 2015; Perry et al., 2015; Ray and Maunsell, 2010; van Pelt et al., 2018) while gamma power undergoes nonlinear changes (Jia et al., 2013; Orekhova et al., 2018b; Salelkar et al., 2018).

Local field potential (LFP) studies of the monkey primary visual cortex have shown that the increase in gamma frequency caused by increase in either luminance contrast (Hadjipapas et al., 2015; Henrie and Shapley, 2005) or temporal frequency of moving gratings (Salelkar et al., 2018) was paralleled by a rise in neuronal spiking rate. This suggests that the increase of gamma frequency is associated with growing neural excitation in V1, and particularly with excitation of inhibitory circuitry that have been shown to play the major role in regulating the frequency of gamma oscillations (Anver et al., 2011; Ferando and Mody, 2013; Mann and Mody, 2010).

The same changes in the stimulation properties that induce a linear increase in gamma frequency lead to nonlinear — often bell-shaped — changes in gamma power in monkey LFP (Hadjipapas et al., 2015; Jia et al., 2011; Jia et al., 2013; Lowet et al., 2015; Salelkar et al., 2018). In a recent MEG study (Orekhova et al., 2018b), we applied high-contrast gratings drifting at different rates (motion velocities) and found the same bell-shaped changes in the gamma response power as those described in the LFP studies on monkeys. The majority of our healthy subjects increased visual GR power from the static to the slowly moving (1.2°/s) stimulus and then suppressed the response at higher velocities of 3.6 °/s and 6.0 °/s. These findings are consistent with those of van Pelt and co-authors (van Pelt et al., 2018), who found a continuous growth in the strength of visual gamma response while changing grating velocities from 0°/s to 0.33°/s and further to 0.66 °/s. Given that the fastest velocity used in their study was still below 1.2°/s, the combined results from both studies are consistent with our previous observation that the effect of stimulus velocity on induced gamma power changed from a facilitative to a suppressive one at a stimulus speed of about 1.2°/s (‘suppression transition velocity’) (Orekhova et al., 2018b).

Seemingly paradoxically, the nonlinear velocity-related gamma power changes may be at least partly explained by rising neural excitation — in particular, the tonic excitation of I-neurons. According to modeling studies, the excitation of I-neurons beyond some critical ‘suppression transition’ point results in their loss of synchrony with each other and excitatory pyramidal cells that, in turn, leads to suppression of gamma oscillations (Borgers and Kopell, 2005; Cannon et al., 2014; Kopell et al., 2010). Thus, the gamma suppression at high velocities/temporal-frequencies of visual motion might be explained by this ‘excitatory drive hypothesis’.

If true, the link between strong inhibition and gamma suppression may have important implications for studying the E and I processes in the healthy and diseased human brain. In particular, individual variations in gamma suppression may reflect the capacity of I circuitry to suppress excessive excitation in the network, thus preventing sensory overload. However, the question remained to be answered – whether the observed nonlinear behavior of the MEG gamma oscillations is related to changes in the strength of excitatory drive or to a particular property of the visual stimulation. Indeed, rather than being a consequence of the strong excitatory drive to the visual cortex, the suppression of gamma response power could result from tuning of the V1 neurons to the particular velocity/temporal frequency. Animal studies show that, while cells in the LGN respond to visual stimuli of rather high temporal frequencies, the neurons in the primary visual cortex show band-pass properties (Priebe et al., 2006; Van Hooser et al., 2013). One therefore cannot exclude that a presence of an ‘optimal velocity’ that induces the maximal visual GR power simply reflects the number of V1 neurons ‘tuned’ to a respective velocity/temporal frequency.

Herein, and in order to test these two alternative explanations, we modulated the GR by changing the velocity of drifting gratings (stationary and moving at 1.2, 3.6 and 6.0°/s that corresponded to temporal frequencies of 0, 2, 6 and 10 Hz) for two luminance contrast values: 100% (high contrast) and 50% (low contrast).

In terms of the excitatory drive model, concomitant increases in contrast and velocity of the stimulation should have an additive effect. If the transition from enhancement to suppression of the GR requires a certain level of excitatory drive, then this level will be reached at a relatively higher motion velocity in case of lower contrast. The excitatory drive model therefore predicts that the ‘gamma suppression transition velocity’ would be higher at the 50%, as compared to the 100%, contrast.

The velocity-tuning model predicts a different outcome from that of the excitatory drive model. In this case, lowering contrast from 100% to 50% either will not affect the ‘transition point’ or shift it to *lower* velocity. The shift to a lower velocity in this case can be expected, because a decrease in the stimulus contrast shifts the tuning of the motion-sensitive V1 neurons to lower temporal frequencies (Alitto and Usrey, 2004; Livingstone and Conway, 2007; Priebe et al., 2006).

Similar to gamma, the oscillations within the alpha-beta range are also thought to reflect the level of cortical activation (Romei et al., 2008; Yuan et al., 2010). Whereas suppression of the alpha-beta oscillations has repeatedly been shown to accompany an active state of cortical networks related to processing of incoming sensory information, the ‘high-alpha’ states are associated with cortical idling or active inhibition of task-irrelevant processes that might otherwise interfere with task performance (Palva and Palva, 2011; Pfurtscheller and da Silva, 1999). Using the visual motion paradigm similar to that used in our study, Scheeringa et al have shown that the power of alpha-beta and gamma oscillations was inversely proportional to changes in the blood oxygen level-dependent (BOLD) signal in the visual cortex, although at different cortical depths (Scheeringa et al., 2016). However, the powers of alpha-beta and gamma oscillations are not interchangeable indexes of cortical activation. In particular, changes in the velocity of visual motion affected alpha and gamma oscillations differently in the LFP in monkeys (Salelkar et al., 2018). There is evidence that visual gamma mainly reflects feed-forward processes (Roberts et al., 2013; van Kerkoerle et al., 2014), while alpha and beta power in the visual cortex are strongly modulated by top-down influences from upstream cortical areas (Liu et al., 2016; Snyder and Foxe, 2010). Therefore, we expected that the bell-shaped pattern of power changes would be specific for the MEG gamma band and would not be mirrored by opposing changes in alpha-beta activity (i.e. gamma response suppression – alpha facilitation).

Overall, our current MEG study pursued these two main goals. Firstly, we tested whether the strength of excitatory drive is the main factor contributing to the initially facilitative, and then suppressive, effects of velocity on the visual gamma response power. Secondly, by comparing the effects of increasing excitatory drive on gamma and alpha oscillations, we sought to unravel the unique information regarding activation of the visual that is conveyed by each of the these two frequency bands.

## Methods

### Participants

Seventeen neurologically healthy subjects (age 18-39, mean=27.2, sd=6.1; 7 males) were recruited to the study. All participants had normal or corrected to normal vision. The informed consent form was obtained from all the participants. The study has been approved by the ethical committee of MSUPE.

### Experimental task

The visual stimuli were generated using Presentation software (Neurobehavioral Systems Inc., USA). We used a PT-D7700E-K DLP projector to present images with a 1280 x 1024 screen resolution and a 60 Hz refresh rate. The experimental paradigm is schematically presented in figure 1. The stimuli were gray-scale sinusoidally modulated annular gratings presented at ∼100% or 50% of Michelson contrast. Luminance of the display, measured from the eye position, was 46 Lux during presentation of both 50% and 100% contrast stimuli and 2 Lux during inter-stimulus intervals. The gratings had a spatial frequency of 1.66 cycles per degree of visual angle and covered 18 × 18 degrees of visual angle. They appeared in the center of the screen over a black background and drifted to the center with one of four velocities: 0, 1.2, 3.6, or 6.0°/s, (0, 2, 6 and 10 Hz temporal frequency) referred to as ‘static’, ‘slow’, ‘medium’, and ‘fast’. Each trial began with 1200 ms presentation of a fixation cross in the center of the display over a black background that was followed by the presentation of the grating that either remained static or drifted with one of the three velocities. After a randomly selected period of 1200–3000 ms, the movement stopped or the static stimulus disappeared. To keep participants alert, we asked them to respond to the change in the stimulation flow (stop of the motion or disappearance of the static stimulus) with a button press. If no response occurred within 1 second, a discouraging message, “too late!” appeared and remained on the screen for 2000 ms, after which a new trial began. The static stimulus and those drifting at different rates were intermixed and appeared in a random order within each of three experimental blocks. The participants responded with either the right or the left hand in a sequence that was counterbalanced between blocks and participants. Each of the stimuli types was presented 30 times within each experimental block. In order to minimize visual fatigue and boredom, short (3–6 s) animated cartoon characters were presented between every 2–5 stimuli. The high (100%) and low (50%) contrast gratings were presented in different experimental sessions and the order of these sessions was counterbalanced between subjects.

**Fig. 1.**
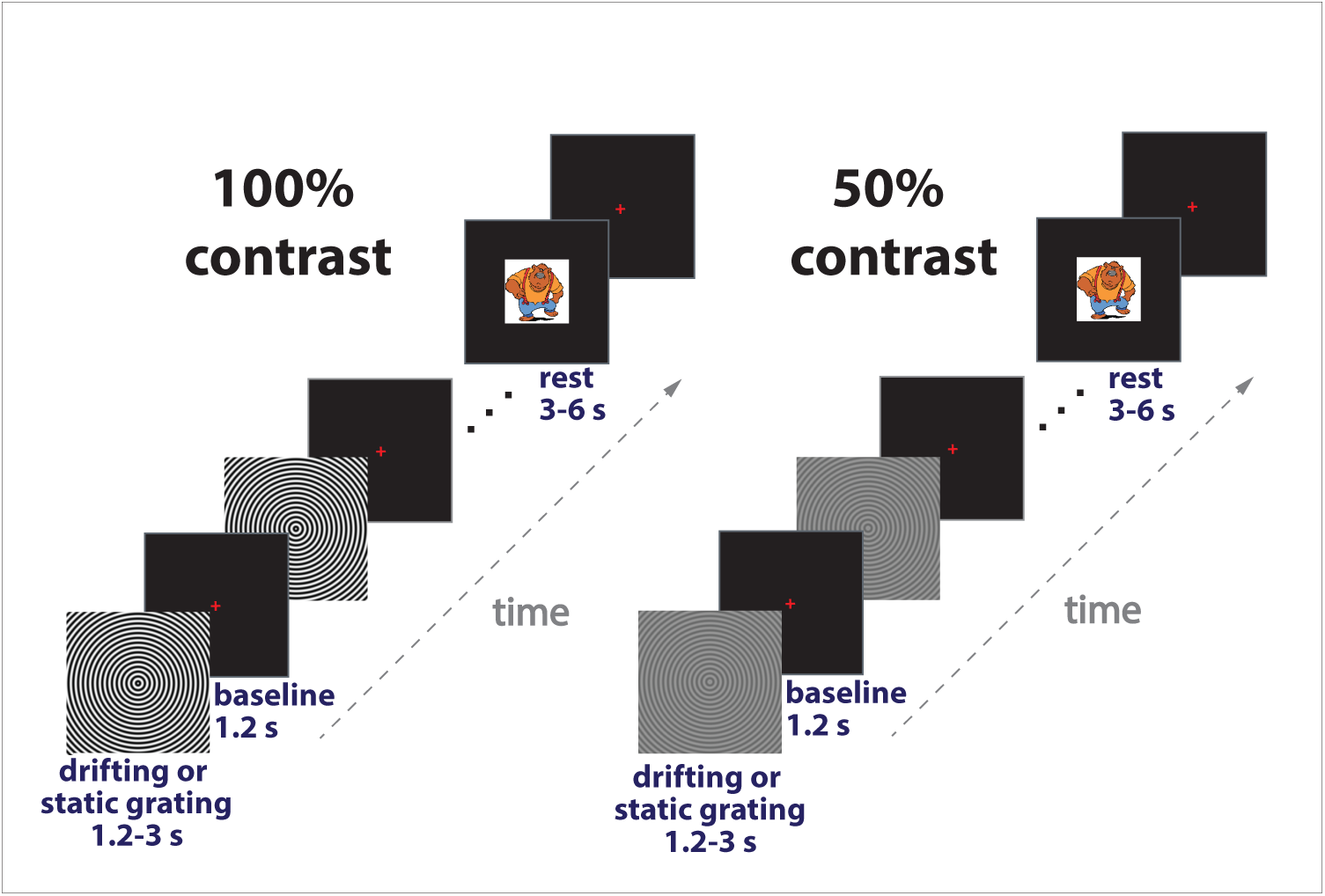
Experimental design. Each trial began with the presentation of a fixation cross that was followed by an annular grating that remained static (0°/s) or drifted inward for 1.2-3 s at one of the three velocities: 1.2, 3.6, 6.0°/s. The 100% and 50% contrast gratings were presented in separate sessions. Participants responded to the change in the stimulation flow (disappearance of the static grating or termination of motion) with a button press. Short animated cartoon characters were presented randomly between every 2-5 stimuli to maintain vigilance and reduce visual fatigue.

### Data recording

Neuromagnetic brain activity was recorded using a 306-channel detector array (Vectorview; Neuromag, Helsinki, Finland). The subjects’ head positions were continuously monitored during MEG recordings. Four electro-oculogram (EOG) electrodes were used to record horizontal and vertical eye movements. EOG electrodes were placed at the outer canti of the eyes and above and below the left eye. To monitor the heartbeats, one electrocardiogram (ECG) electrode was placed at the manubrium sterni and the other one at the mid-axillary line (V6 ECG lead). MEG, EOG, and ECG signals were recorded with a band-pass filter of 0.03–330 Hz, digitized at 1 000 Hz, and stored for off-line analysis.

### MEG data preprocessing

The data was first de-noised using the Temporal Signal-Space Separation (tSSS) method (Taulu and Hari, 2009) implemented in MaxFilter™ (v2.2) with parameters: ‘-st’=4 and ‘-corr’=0.90. For all three experimental blocks, the head origin position was adjusted to a common standard position. For further pre-processing, we used the MNE-python toolbox (Gramfort et al., 2013) as well as custom Python and MATLAB^®^ (TheMathWorks, Natick, MA) scripts.

The de-noised data was filtered between 1 and 200 Hz and down sampled to 500 Hz. To remove biological artifacts (blinks, heart beats, and in some cases myogenic activity), we then applied independent component analysis (ICA). The MEG periods with too high (4e-10 fT/cm for gradiometers and 4e-12 fT for magnetometers) or too low (1e-13 fT/cm for gradiometers and 1e-13 fT for magnetometers) amplitudes were excluded from the analysis. The number of independent components was set to the dimensionality of the raw ‘SSS-processed’ data (usually around 70). We further used an automated MNE-python procedure to detect EOG and ECG components, which we complemented with visual inspection of the ICA results. The number of rejected artifact components was usually 1-2 for vertical eye movements, 0-3 for cardiac, and 0-6 for myogenic artifacts.

The ICA-corrected data was then filtered with a discrete Fourier transform filter to remove power-line noise (50 and 100 Hz) and epoched from −1 to 1.2 sec relative to the stimulus onset. We then performed time-frequency multitaper analysis (2.5 Hz step; number of cycles = frequency/2.5) and excluded epochs contaminated by strong muscle artifacts via thresholding the high-frequency (70 - 122.5 Hz) power. For each epoch, the 70 - 122.5 Hz power was averaged over sensors and time points and the threshold was set at 3 standard deviations of this value. The epochs left were visually inspected for the presence of undetected high-amplitude bursts of myogenic activity and those contaminated by such artifacts were manually marked and excluded from the analysis. After rejection of artifacts the average number of epochs for the 100% contrast condition was 79, 78, 78, and 79 for the ‘static’, ‘slow’, ‘medium’ and ‘fast’ conditions, respectively. For the 50% contrast, the respective values were 79, 80, 80, and 82.

### Structural MRI

Structural brain MRIs (1 mm^3^ T1-weighted) were obtained for all participants and used for source reconstruction.

### Time-frequency analysis of the MEG data

The following steps of the data analyses were performed using Fieldtrip Toolbox functions (http://fieldtrip.fcdonders.nl; (Oostenveld et al., 2011)) and custom scripts developed within MATLAB.

We subtracted the averaged evoked responses from each data epoch. This decreases the contribution of phase-locked activity that could be related to the appearance of the visual stimulus on the screen and the effect of photic driving related to the temporal frequency of the stimulation, screen refresh rate, or their interaction.

The lead field was calculated using individual ‘single shell’ head models and cubic 6 mm-spaced grids linearly warped to the MNI-atlas-based template grid. The time-frequency analysis of the MEG data was then performed in the following two steps.

*First,* we tested for the presence of reliable clusters of gamma facilitation and alpha suppression at the source level using the DICS inverse-solution algorithm (Gross et al., 2001) and the cluster-based permutation test (Nichols and Holmes, 2002). The frequency of interest was defined individually for each subject and condition based on the sensor data (only the data from gradiometers were used at this stage). For the gamma range, we performed time-frequency decomposition of pre-stimulus (‘pre’: −0.9 to 0 s) and post-stimulus (‘post’: 0.3 to 1,2 s) MEG signals using 8 discrete prolate spheroidal sequences (DPSS) tapers with ±5 Hz spectral smoothing. For the low frequency range, we used 2 DPSS tapers and ±2 Hz spectral smoothing. We then calculated the average (post-pre)/pre ratio and found the 4 posterior sensors with the maximal post-stimulus increase in gamma power (45-90 Hz) and those with the maximal post-stimulus power decrease in the low-frequency band (7-15 Hz). By averaging those 4 channels, we have found the frequencies corresponding to the maximal post-stimulus power increase (for gamma) or decrease (for low frequency range). The frequency ranges of interest were then established within 35-110 Hz and 5-20 Hz limits, where the ratios exceeded 2/3 of the respective peak value. The center of gravity of the power over these frequency ranges was used as the peak frequency of the gamma and the slow responses, respectively. The time-frequency decomposition was then repeated while centered at these peak frequencies. Common source analysis filters were derived for combined pre- and post-stimulus intervals using DICS beamforming with a 5% lambda parameter and fixed dipole orientation (i.e, only the largest of the three dipole directions per spatial filter was kept). The filter was then applied separately to the pre- and post-stimulus signals. Subsequently, we calculated univariate probabilities of pre- to post-stimulus differences in single trial power for each voxel using T-statistics. We then performed bootstrap resampling (with 10 000 Monte Carlo repetitions) between ‘pre’ and ‘post’ time windows to determine individual participants’ maximal source statistics based on the sum of T-values in each cluster.

*Second*, in order to analyze the response parameters at the source maximum, we used linearly constrained minimum variance (LCMV) beamformers (VanVeen et al., 1997). The spatial filters were computed based on the covariance matrix obtained from the whole epoch and for the three experimental conditions with a lambda parameter of 5%. Prior to beamforming, the MEG signal was band-passed between 30 and 120 Hz for the gamma-range and low-passed below 40 Hz for the low frequency range. The ‘virtual sensors’ time series were extracted for each brain voxel and time-frequency analysis with DPSS multitapers (∼1 Hz frequency resolution) was performed on the virtual sensor signals. For the gamma range, we used ±5 Hz smoothing. For the low-frequency range, this parameter was ±2 Hz. The ‘*maximally induced*’ gamma voxel was defined as the voxel with the highest relative post-stimulus increase of 45-90 Hz power in the ‘slow’ motion velocity condition within the visual cortical areas (i.e. L/R cuneus, lingual, occipital superior, middle occipital, inferior occipital, and calcarine areas according to the AAL atlas (Tzourio-Mazoyer et al., 2002)). The ‘slow’ condition was selected because reliability of the gamma response was highest in this condition. The ‘*maximally suppressed*’ voxel in the low-frequency range was then defined as the voxel with the strongest relative suppression of 7-15 Hz power. The weighted peak parameters for the gamma and the low-frequency activity were calculated for the average spectra of the virtual sensors in 25 voxels closest to, and including, the ‘maximally induced’ and ‘maximally suppressed’ voxels, respectively, using the approach described for the sensor level analysis. For each individual and condition, we also assessed the reliability of the pre- to post-stimulus power changes at these 25 averaged voxels. The change was considered reliable if its absolute peak value was significant with p<0.0001 (Wilcoxon signed rank test). Coordinates of the voxels demonstrating greatest power changes were defined in MNI coordinates (Supplementary Tables).

## Results

We measured MEG brain responses induced by concentric visual grating drifting inwardly at four velocities (0, 1.2, 3.6, 6.0 °/s) and two contrast levels (50% and 100%). Below we present the results separately for the gamma and the alpha-beta ranges.

### Gamma frequency range

#### Localization and reliability

Figure 2 shows grand average source localization of the induced gamma responses.

**Fig. 2.**
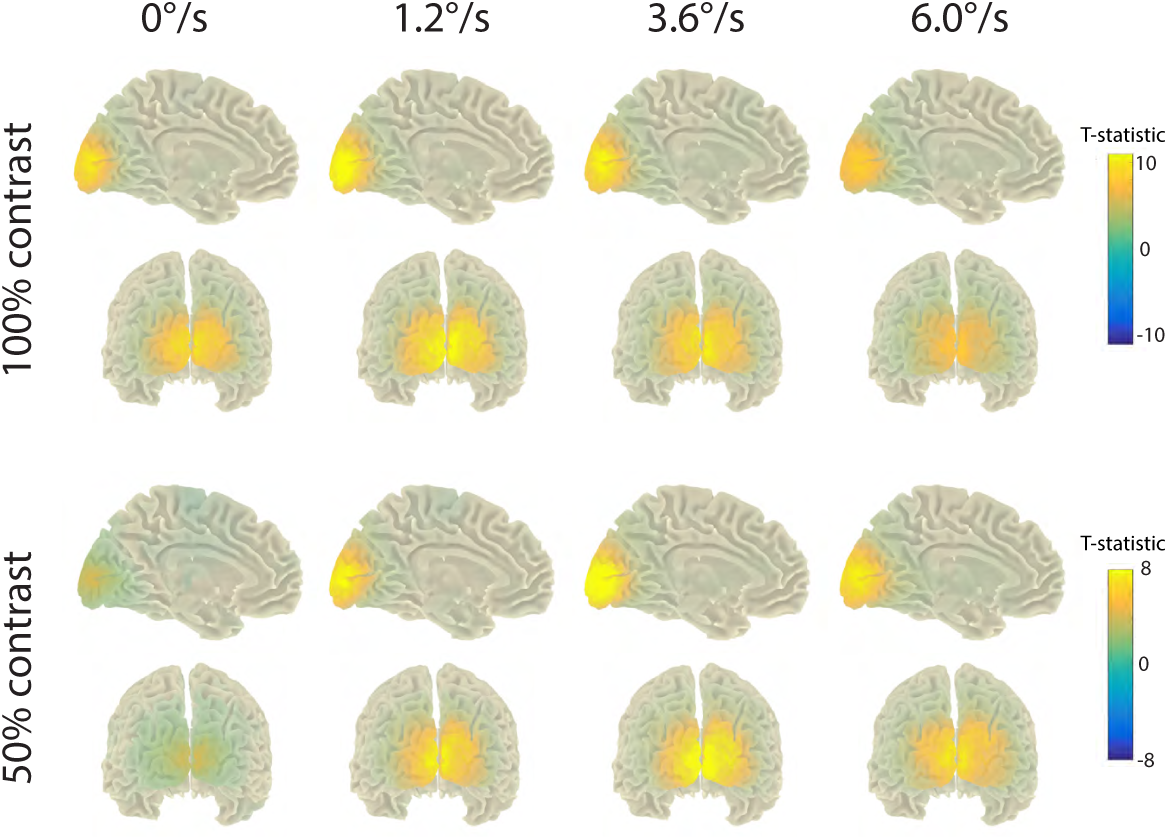
Grand average statistical maps of the cortical GRs to drifting visual gratings. The maps are given for the weighted peak gamma power, separately for the two luminance contrasts and four motion velocities. Positive sign of the T-statistics corresponds to stimulation-related increase in gamma power. Note different scales for the 50% and 100% contrasts and that the magnitude of the gamma response was affected by both contrast and velocity.

For the 100% contrast stimuli, significant (p<0.05) activation clusters with maxima in visual cortical areas were found in all participants and motion velocity conditions. For the 50% contrast, the significant activation clusters were absent in one participant in all velocity conditions, in two participants in the static condition, and in one participant in the 6.0°/s condition.

The grand average and individual gamma response spectra at the ‘maximally induced’ group of voxels are shown in figures 3 and 4. The participants A007, A008, and A016 had no reliable increase in gamma power (p>0.0001, see Methods for details) at 50% contrast in at least one of the velocity conditions (A007: all velocities; A008 and A016: 0°/s). In all other participants, the increases in gamma power at the selected voxels were significant according to our criteria in all conditions.

**Fig. 3.**
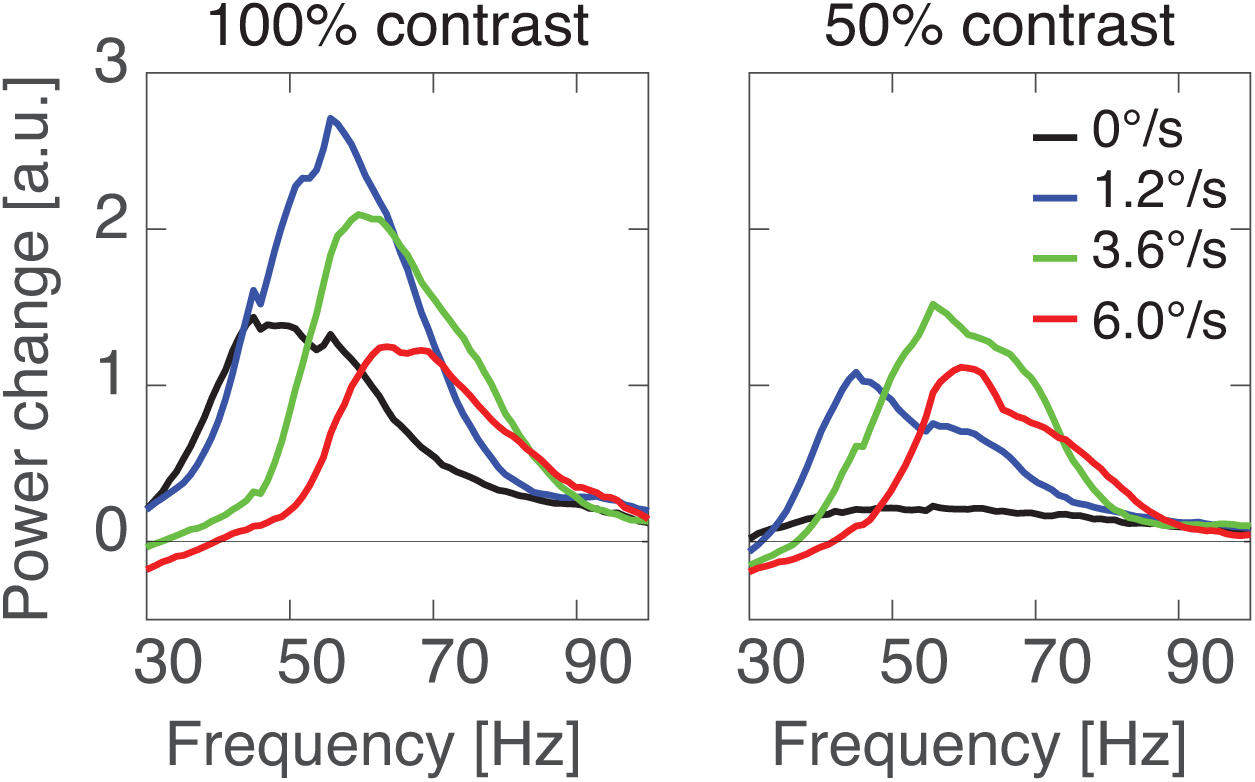
Grand average spectra of cortical GRs to the gratings of 100% (left panel) and 50% (right panel) contrasts drifting at four motion velocities. Here and hereafter, the GR power is calculated as (post-pre)/pre ratio, where ‘pre’ and ‘post’ are the spectral power values in the −0.9 to 0 and 0.3 to 1.2 s time windows relative to the stimulus onset.

The peaks that were unreliable according to at least one of the criteria (i.e. either the absence of an activated cluster or low probability (p>0.0001) of gamma increase at the selected voxels) were further excluded from analyses of the peak frequency and the position of the maximally induced voxel. Note that exclusion of the unreliable gamma peaks affected the degrees of freedom in the corresponding ANOVAs presented below.

In the majority of cases, the voxel with the maximal increase in gamma power was located in the calcarine sulcus (see the Supplementary Table 1A and 1B for coordinates of the ‘maximally induced’ voxel). There was a significant effect of Velocity for the ‘z’ coordinates (F_(3,39)_=4.7, e=0.84, p<0.05). In the ‘fast’ condition, the voxel with the maximal increase in gamma power was located on average 2.6 mm more superior relative to the other 3 conditions (Z coordinate for the static: 0.61 cm, slow: 0.54 cm, medium: 0.52 cm, fast: 0.82 cm). There was also Velocity*Contrast interaction for the ‘y’ coordinate (F_(3,39)_=3.6, e=0.59, p<0.05), that is explained by a relatively more posterior source location of the gamma maximum in the ‘fast’ low-contrast condition than in the ‘fast’ high-contrast condition (−9.5 vs −9.2 cm). Considering the small condition-related differences in the position of the voxel with the maximal increase in gamma power (<0.6 cm, which was the size of the voxel used for the source analysis), all the stimuli activated largely overlapping parts of the primary visual cortex.

#### Effects of contrast and velocity

##### Frequency

Figure 5A shows group mean gamma peak frequency values for the two contrast and four velocity conditions. Gamma frequency was strongly affected by motion velocity of the grating (F_(3,39)_=78.2,e=0.62 p<1e-6) and, to a lesser extent, by its contrast (F_(1,13)_=14.3, p<0.01). Inspection of figure 5A shows that increase in velocity resulted in substantial increase of gamma frequency (∼12-17 Hz). For the full-contrast grating the frequency increased from 51.6 Hz in case of the static to 68.6 Hz in case of the fast moving stimulus. For the 50% contrast the respective values were 51.1 and 63.0 Hz. Increasing contrast, on the other hand, led to 5Hz increase in frequency of gamma response to moving stimuli.

**Fig. 4.**
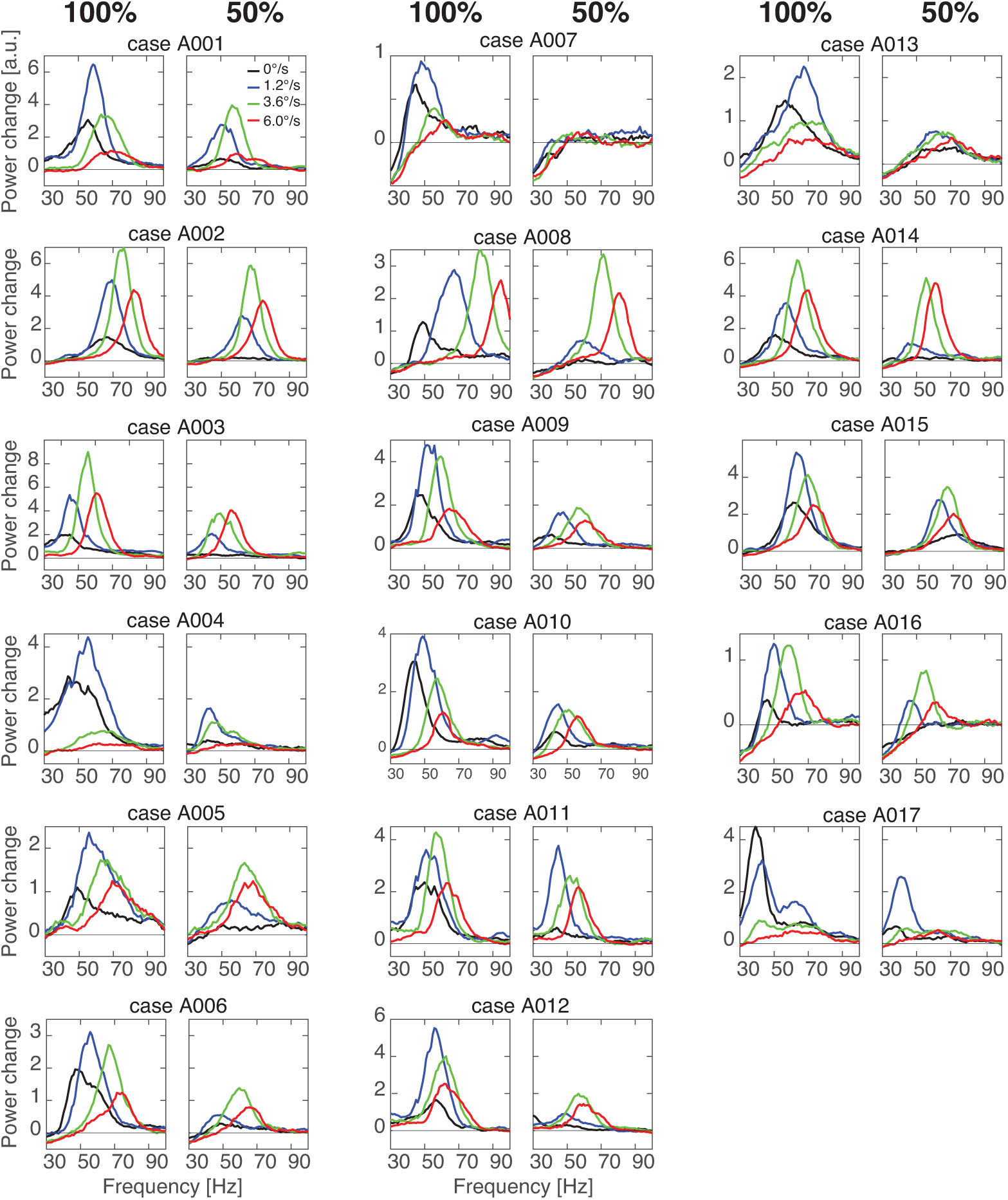
Individual spectra of cortical GRs. The plots are the same as in Fig. 3, see Methods for further details.

**Fig. 5.**
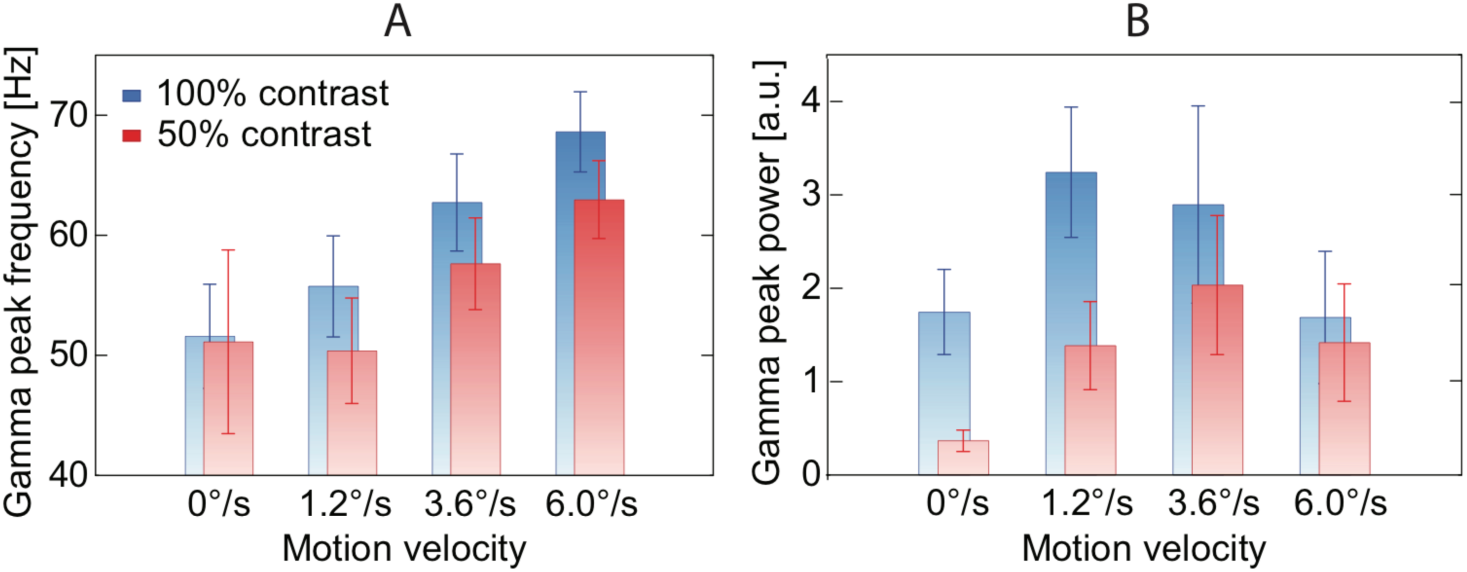
Grand average peak frequency (A) and power (B) of cortical GRs as a function of Contrast and motion Velocity. Vertical bars denote 0.95 confidence intervals.

##### Power

Figure 5B shows group mean GR peak power values for the two contrast and four velocity conditions. There was highly significant effect of Contrast (F_(1,16)_=66.2,p<1e-6) on GR peak power, which is explained by a generally higher gamma power in case of high, as compared to low contrast. The significant effect of Velocity (F_(3,48)_=9.3, e=0.46, p<0.01) is due to an initial increase in GR power from the static to slow condition (F_(1,16)_=43.1, p<1e-5) followed by a decrease from the medium to the fast velocity condition (F_(1,16)_=14.2, p<1e-4). Importantly, there was highly significant Contrast*Velocity interaction effect (F_(3,48)_=12.4, e=0.78, p<1e-4). The contrast had a stronger effect on GR power for the static and slowly moving stimuli, as compared to the faster (medium and fast) velocities. As a result, the maximum of the inverted bell-shaped power vs. motion velocity curve moved to higher velocity in the low contrast condition, as compared to the full contrast one.

Generally, the results confirmed our previous finding of an inverted bell-shaped dependency of GR power on the velocity of visual motion and extended them to two contrast conditions. The significant Contrast*Velocity interaction showed that the form of this bell-shaped curve was modulated by the luminance contrast in such a way that the transition to GR suppression at low contrast occurred at a higher velocity, as predicted by the excitatory drive model.

#### Suppression transition velocity

We further sought to investigate whether the shift to higher velocities in the GR suppression transition point under low contrast is a robust phenomenon, which can be detected at the individual subject level. For each subject, we calculated the ‘suppression transition velocity’ (STVel) as the centre of gravity visual motion velocity:

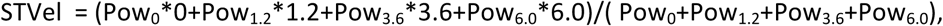

where ‘Pow’ is the weighted GR peak power in the respective velocity condition (i.e. 0, 1.2, 3.6 or 6.0 °/s). Although the STVel may not precisely correspond to the velocity condition for which the GR was at maximum, it reflects the distribution of power between velocity conditions. More importantly, it allows us to characterize contrast-dependent shifts in a more rigorous manner than relying exclusively on the discrete values of motion velocities for which the GR was maximal. Decreasing luminance contrast resulted in a reliable increase in STVel, and thus the GR suppression transition point, in all 17 participants (F_(1,16)_=202.0; p<1e-6; Fig. 5A).

Lowering contrast led to slowing of gamma oscillations (Fig. 4A); we therefore also tested if, irrespective of contrast, the point of transition to GR suppression corresponds to a certain gamma frequency. To do this, we approximated the suppression transition frequency (STFreq, i.e. the gamma peak frequency that corresponds to the maximal gamma response) as the center of gravity of the GR power for each subject (i.e., in the same way as we did for the STVel):

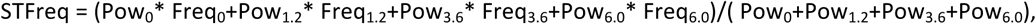

where ‘Freq’ and ‘Pow’ are the peak gamma response frequency and power in the respective velocity conditions.

The suppression transition frequency was slightly, but significantly, lower for the 50% than for the 100% contrast (60.5 Hz vs 63.1 Hz; F_(1, 13)_=10.2, p<0.01).

### Rank-order consistency of gamma parameters

Considering the drastic stimulation-related changes in gamma parameters, their intra-individual stability across experimental conditions (i.e. ‘rank-order consistency’) remained an open question. In order to investigate if the power and frequency of GR measured in one experimental condition predicts the subject’s rank position in another experimental condition, we calculated two types of correlations: 1) between contrasts for the respective velocity conditions and 2) between velocities, separately for each of the contrasts. Since frequency, but not power, of the visual gamma oscillations is strongly affected by age in adults (Gaetz et al., 2012; Orekhova et al., 2018b), we calculated partial correlations taking age as a nuisance variable with frequency.

The correlations between contrast conditions were generally very high for gamma frequency (all R’s>0.89; all p’s <0.0001), with the noticeable exception of the static stimulus (R=0.36, n.s.). For power, all the between-contrast correlations were modestly to highly reliable (Fig. 7A). The correlation between STVel values measured at the 50% and 100% contrast was high (Fig. 6B, STVel: R_(17)_=0.91, p<1e-6), suggesting that this measure reliably characterises a subject’s rank position in the group, irrespective of contrast changes.

**Fig. 6.**
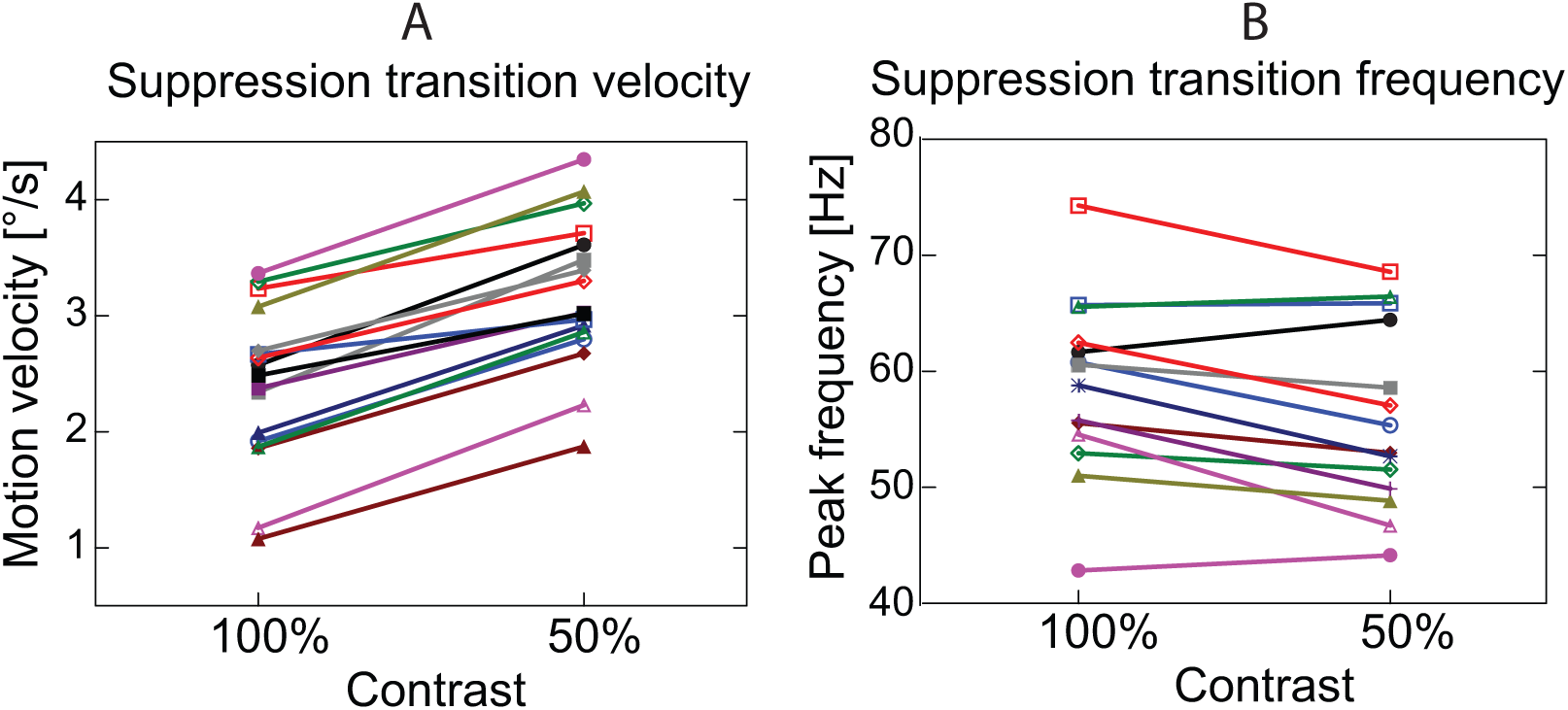
Effect of contrast on the gamma suppression transition point: individual variability. A. The centre of gravity of visual motion velocity that approximates the GR suppression transition point (‘suppression transition velocity’ - STVel). In every single subject, the STVel increased with decreasing contrast. B. The centre of gravity of gamma peak frequency that approximates the peak frequency of the GR at the gamma suppression transition point (‘suppression transition frequency’ - STFreq). The STFreq slightly, but significantly, decreased with decreasing the contrast.

**Fig. 7.**
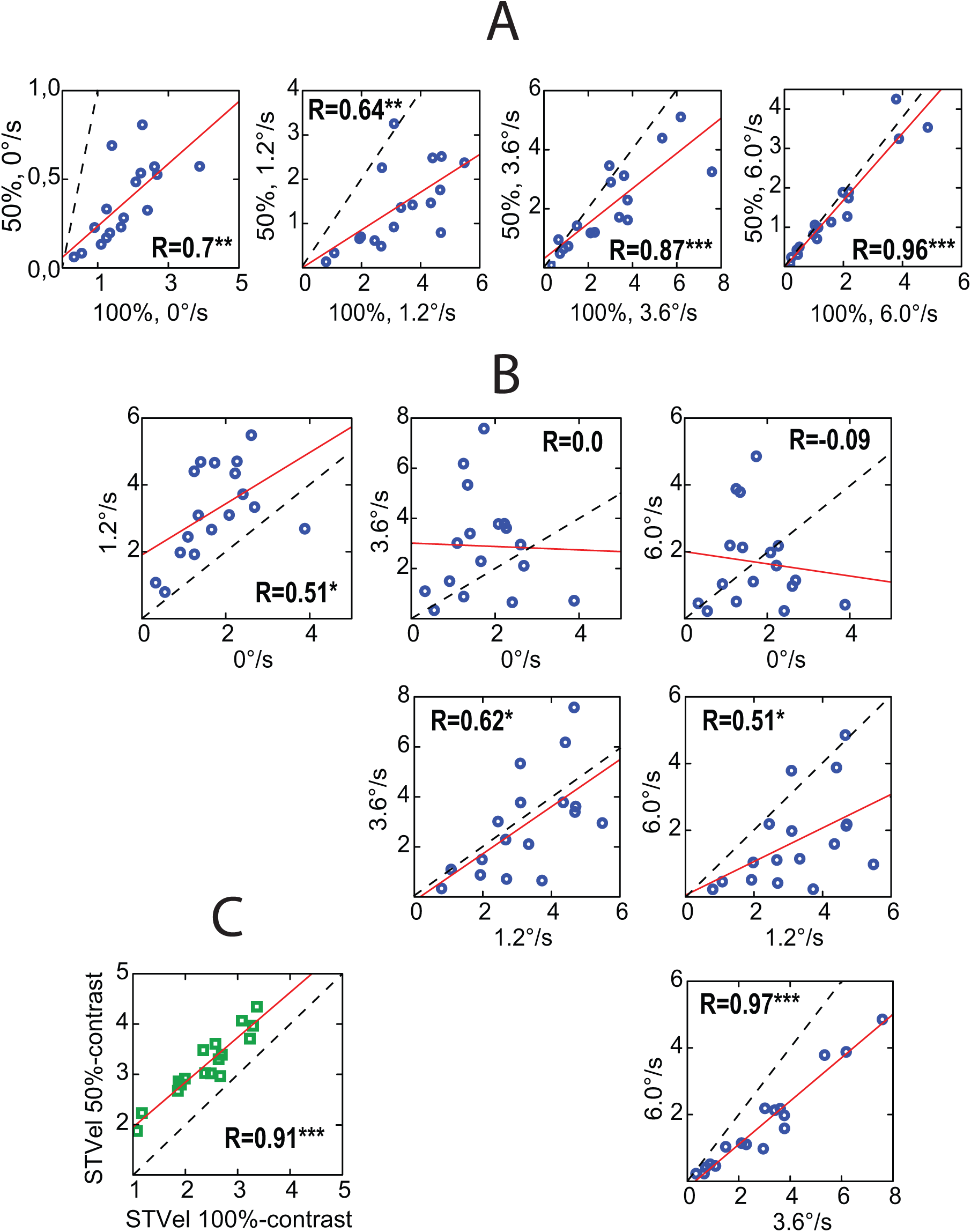
Rank-order consistency of the GR power across conditions. A. Correlations between GR power measured at 50% and 100% contrasts. B. Correlations between GR power values measured in different velocity conditions at the 100% contrast (see Supplementary Figure 1 for a similar figure for the 50% contrast stimuli). Blue dots in A and B denote individual gamma response power values. C. Correlations between STVel values at the two contrasts. Green squares denote individual STVel values. The linear regression is shown in red. Dashed lines in all plots correspond to the axis of symmetry.

The correlations between velocity conditions were also generally high for frequency (Supplementary Table 3), but varied in case of power. The power of the GR elicited by static stimuli did not predict the power of the GR elicited by gratings moving with ‘medium’ of ‘fast’ velocities (Fig. 7B). The results were very similar for the two contrasts (Supplementary Figure 1).

To sum up, rank-order consistency for gamma frequency was generally high across all conditions except for stationary stimuli. For gamma power, a rank-order consistency between stationary and medium-to-fast moving gratings was lacking.

### Alpha-beta frequency range

#### Localization and reliability

Figure 8 shows grand average source localization of the alpha-beta response to the moving visual gratings.

**Fig. 8.**
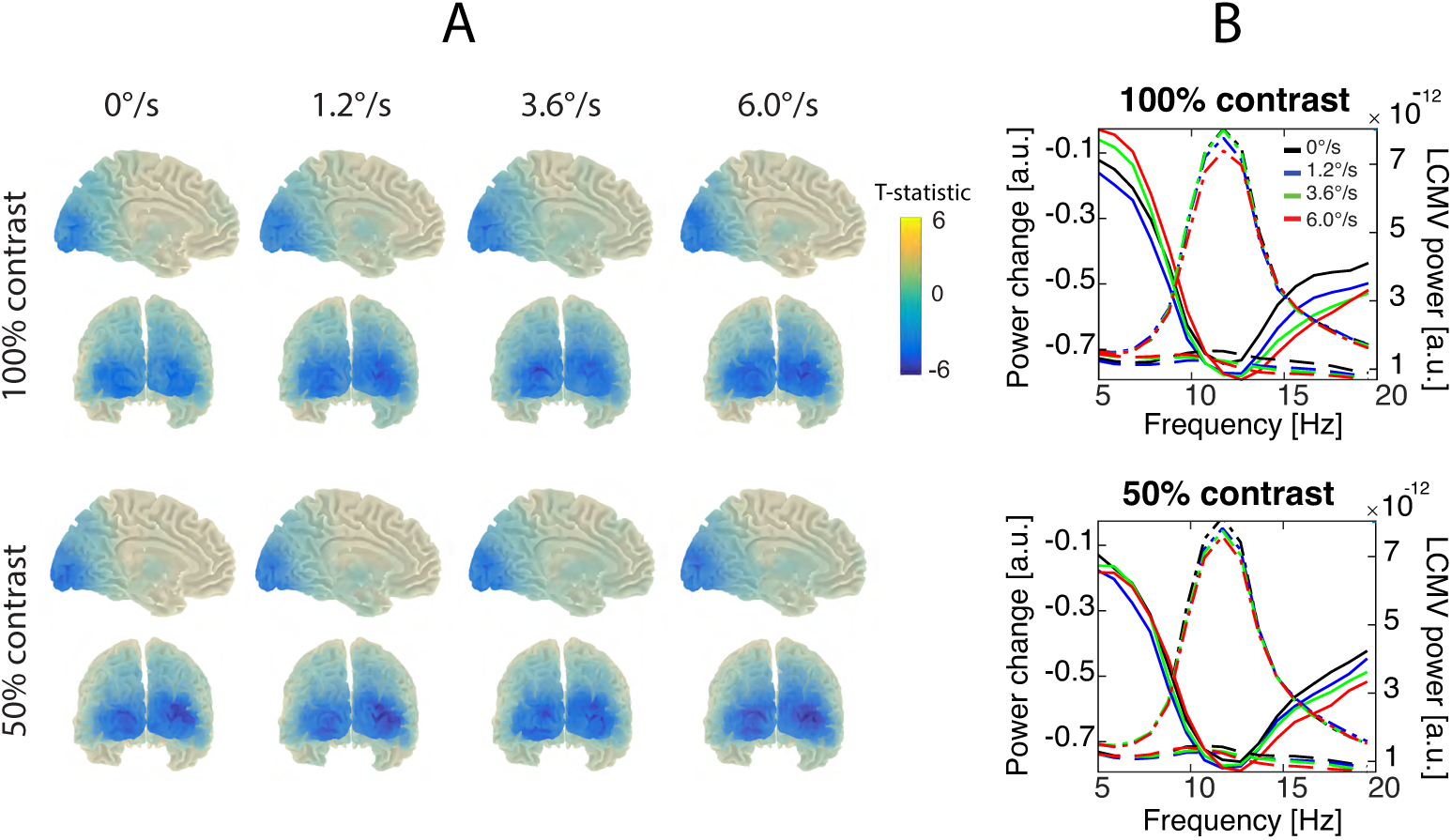
Grand average power changes in the alpha-beta range. A. Statistical maps of the stimulation-related changes in the two luminance contrasts and four motion velocity conditions. Blue color corresponds to suppression of the alpha-beta power. B. Spectral changes at a selection of ‘maximally suppressed’ voxels. Dashed lines show absolute power spectra (right y-axis, arbitrary units) corresponding to the periods of visual stimulation with moving gratings. The dot -dashed lines of the same colors show the power spectra for the respective pre-stimulus intervals (also right y-axis). Solid lines show respective stimulation-related changes in power ([post-pre]/pre ratio, left y-axis, arbitrary units). Neither contrast nor velocity had significant effects on the magnitude of the alpha-beta suppression.

Clusters of significant (p<0.05) alpha-beta power suppression were found in nearly all subjects and conditions. For the 100% contrast stimuli, the clusters were absent in one subject in all velocity conditions and in on subject in the static condition. The suppression measured at the selection of the ‘maximally suppressed’ voxels was significant in all motion velocity conditions at 100% contrast. For the 50% contrast stimuli, a significant cluster for alpha-beta suppression was absent in only one subject and only in response to the static stimulus. In yet another subject, the alpha-beta suppression measured at the voxels’ selection was not significant (p>0.0001) for the 1.2 °/s condition. The grand average spectra in the alpha-beta range for the maximally suppressed voxels are shown in figure 8.

ANOVA with factors Contrast and Velocity revealed neither significant main effects of Contrast and Velocity nor Contrast*Velocity interaction for the ‘x’, ‘y’, or ‘z’ coordinates of the ‘maximally suppressed’ voxel (all p’s>0.15).

We then used ANOVA with factors Band, Contrast, and Velocity to compare positions of the voxels with maximal alpha-beta suppression and those with maximal gamma increase. The effect of Band was highly significant for the absolute value of the ‘x’ coordinate (F_(1,16)_=81.9, p<1e-6). The maximally suppressed alpha voxel was positioned substantially more laterally (on average 25 mm from the midline) than that of the maximally induced gamma voxel (on average 8 mm from the midline). As can be seen from Supplementary Table 2(A,B), the maximally suppressed alpha voxel was more frequently located in the lateral surface of the occipital lobe than in calcarine sulcus or cuneus region. No Band related differences in the y or z coordinates were found.

#### Effects of contrast and velocity

To check if the magnitude or peak frequency of the alpha-beta suppression response depends on the properties of the stimulation, we performed ANOVA with factors Contrast and Velocity.

##### Power

There was no significant effects of Contrast or Velocity or their interaction effect for the alpha-beta suppression magnitude (all p’s>0.5).

##### Frequency

The peak frequency of the alpha-beta suppression sightly, but significantly, increased with increasing motion velocity (Velocity: F_(3,36)_=7.1, e=0.57, p<0.01; 11.8 Hz for 0°/s, 11.9 Hz for 1.2°/s, 12.0Hz for 3.6°/s, 12.2 Hz for 6.0°/s).

## Discussion

We tested for an additive effect of contrast and velocity of drifting visual gratings on the power of induced MEG gamma oscillations. For both full-contrast and low-contrast gratings, the power of gamma response initially increased with increasing motion velocity and was then suppressed at yet higher velocities. Most importantly, we found that lowering contrast led to a highly reliable shift in the gamma suppression transition point to higher velocities. This finding provides an experimental proof for the crucial role of intensive excitatory input in GR suppression. The systematic relationship between the intensity of the visual input and the strength of the neural response was clear for gamma oscillations and absent in the alpha-beta frequency range.

### The role of excitatory drive in induced gamma response

In line with the previous studies (Jia et al., 2013; Perry et al., 2015; Ray and Maunsell, 2010; Roberts et al., 2013; van Pelt et al., 2018), we found that the peak frequency of the visual gamma response increased with increasing luminance contrast (Fig. 4A). As an increase in luminance contrast causes a rise of neuronal spiking rate (Bartolo et al., 2011; Hadjipapas et al., 2015; Ray and Maunsell, 2010), contrast can be considered as a proxy for excitatory drive (Lowet et al., 2015). Therefore, contrast-dependent changes in gamma frequency and amplitude can be attributed to changes in excitatory input to the visual cortex.

Also in accordance with our previous results—and those of others— (Muthukumaraswamy and Singh, 2013; Orekhova et al., 2015; Orekhova et al., 2018b; Swettenham et al., 2009; van Pelt et al., 2018), we found that the peak frequency of the MEG GR prominently increased (17 Hz on average for the 100% contrast) with increasing motion velocity from 0 to 6 °/s, which corresponds to 0 - 10 Hz of temporal frequency in our study (Fig 4A). No saturation effect has been observed for the gamma frequency even at the highest speeds of stimulation. Most probably, such noticeable acceleration of gamma oscillations at the high stimulus velocity reflects a high excitatory state of the visual cortex caused by fast visual motion. This assumption is supported by studies on monkeys showing that increasing the temporal frequency of drifting gratings from 0 Hz up to approximately 10 Hz leads to a rise in neuronal spiking and an increase in gamma frequency in V1 (Salelkar et al., 2018). Fast visual motion might engage more attention, which may also affect the excitatory state of V1 (Reynolds and Chelazzi, 2004) and thus modulate the gamma frequency during visual motion stimulation (van Pelt et al., 2018). The effect of attention alone is, however, unlikely to explain the observed 12-17 Hz increase in gamma frequency with transition from static to fast-moving stimuli presented herein. Indeed, studies in monkeys have shown that attention has a weak effect on gamma frequency (approximately 3 Hz) (Bosman et al., 2012). Therefore, changes in gamma frequency with increasing motion velocity are most likely accounted for by a rise in bottom-up excitatory input to the visual cortex.

The most important finding of the current study is the effect of luminance contrast on the bell-shaped input-output relationships between visual motion velocity and the magnitude of induced GR. Although at both 50% and 100% contrast, the initially facilitative effect of increasing velocity on gamma response strength was followed by a suppressive one, the maximum of this bell-shaped curve at the 50% contrast shifted to higher velocity, as compared to the 100% contrast (Fig. 3, 5). Inspection of the individual data has shown that with lowering contrast, the approximated velocity of the gamma suppression transition increased in every single subject (Fig. 6A, 7C). This finding suggests that a certain level of excitation is necessary for the gamma suppression to occur, which in case of the low contrast is achieved at a relatively high velocity. Thus, our experimental results strongly support the “excitatory drive” model that predicts an additive effect of increasing contrast and velocity/temporal-frequency on the gamma suppression transition point.

The shift of the GR suppression transition point to higher velocities with lowering luminance contrast contradicts the ‘velocity tuning’ model. Animal studies have shown that at lower contrast, the majority of responsive neurons in V1 (Alitto and Usrey, 2004; Livingstone and Conway, 2007; Priebe et al., 2006), LGN (Alitto and Usrey, 2004), and even the retina (Shapley and Victor, 1978, 1981) are turned to ***lower*** temporal frequencies. Therefore, if the maximal GR would correspond to an ‘optimal’ velocity, the suppression transition point would rather shift to a lower drifting rate when contrast is reduced.

Another and more general argument against the ‘velocity tuning’ hypothesis is the mismatch in velocity tuning of the V1 multiunit activity and that of the V1 gamma response. A recent study in nonhuman primates have shown that the gamma response power starts to suppress at lower temporal frequencies than those eliciting maximal spiking rate in multiunit activity. Specifically, at the temporal frequencies of visual motion corresponding to our ‘medium’ and ‘fast’ velocities, the gamma response already passed its maximum and started to decrease, while neural spiking continued to increase or saturated (see Fig. 4 in (Salelkar et al., 2018)).

Although our findings favor the excitatory drive model, apart from the excitatory drive, there may be other factors that also affect gamma parameters at the suppression transition point. The low-contrast stimuli, as compared with the high-contrast ones, generally induced lower-amplitude gamma responses and triggered the transition to the gamma suppression at lower amplitude and frequency of the GR (Fig. 5A, 6B). Lower contrast stimuli are known to activate fewer neurons and to produce lower spiking rates in neural circuitry involved in gamma synchronization (Henrie and Shapley, 2005). This explains the lower power and frequency of the gamma response at the suppression transition point in case of the lower contrast.

### State and trait dependency of visual gamma parameters

Previous studies show that parameters of visual gamma oscillations are strongly genetically determined (van Pelt et al., 2012) and highly reproducible when measured across time (Hoogenboom et al., 2006; Muthukumaraswamy et al., 2010). As such, they represent constitutional traits that are determined by neurophysiological and neuro-anatomical factors (Schwarzkopf et al., 2012; van Pelt et al., 2018).

While being individually stable in a single experimental condition, the gamma parameters are strongly affected by the properties of the visual stimulation. This raises the question of whether gamma measured in response to different visual stimuli — e.g. those having different contrasts or moving with different speeds — reflect the same or different neurophysiological traits.

Previous studies that sought to find a link between visual gamma oscillations and processes of neural excitation and inhibition in the human brain (Cousijn et al., 2014; Edden et al., 2009; Kujala et al., 2015) measured parameters of the induced gamma response in a single experimental condition. Such an approach implicitly assumes that the features of gamma response do not substantially vary with stimulus properties in term of the rank-order consistency of the individual values. Recently, van Pelt et al (van Pelt et al., 2018) reported high between-condition correlations for gamma peak frequency and power measured at two contrasts (50% and 100%) and at three velocities (0, 0.33°/s, 0.66°/s) that support this view. Importantly, the gamma response power in that study monotonously increased with increasing velocity. This means that all of the stimulation velocities chosen by van Pelt et al were below the ‘gamma suppression transition velocity’ and corresponded to the ascending branch of the bell–shaped curve characterizing the relation between velocity and gamma power.

When we used a broader range of velocities, we found no reliable correlations between the amplitudes of gamma responses caused by static (0°/s) and rapidly moving (3.2°/s, 6.0°/s) stimuli (Fig. 7B). Thus, the intra-individual stability of gamma power across these conditions was lacking, suggesting that the strength of the GR at the ascending and descending branches of the bell-curve is mediated by distinct neural processes. These findings clearly indicate that the analysis of visually-induced changes in gamma oscillations should extend beyond the classical gamma frequency and power metrics in single experimental conditions.

We found a remarkably strong and highly reliable correlation between gamma suppression transition velocities at the 100% and 50% contrasts (Spearman R_(17)_=0.91), pointing to the high within-subject reliability of individual STVel values across different experimental conditions. As we will argue in the following discussion, the STVel is a physiologically meaningful measure that may help to characterize the capacity of the network to regulate a balance between excitation and inhibition.

### Magnitude of the alpha-beta suppression is not sensitive to the strength of visual input

Although stationary and drifting visual gratings lead also to a strong and reliable suppression of the MEG power in the alpha-beta range over the posterior cortical areas (Fig.3), the visually induced alpha-beta response was clearly different from that in the gamma range. *First*, despite apparent overlap (Fig. 2, 8), the spatial localization of the alpha-beta suppression and gamma facilitation significantly differed. Whereas the maximum of the gamma response was localized near the calcarine sulcus, the alpha-beta suppression occurred in more lateral areas of the occipital lobes (see also Supplementary Table 1 and 2 for localization of the maximal effect voxels). This finding is in line with results of previous studies that show more lateral positions of the sources of alpha-beta suppression relative to those of gamma enhancement (Hoogenboom et al., 2006; Koelewijn et al., 2011; Muthukumaraswamy et al., 2016). *Second*, and more importantly, in sharp contrast with the gamma response, neither contrast nor velocity affected power suppression in the alpha-beta frequency range. This lack of a relationship between the intensity of visual stimulation and the degree of the alpha-beta suppression questions the commonly held notion that the magnitude of visual alpha suppression indexes the excitatory state of the visual cortex. On the other hand, our findings do not contradict the numerous results associating relative decreases and increases in visual alpha-band activity with attention-related influences modulating excitability of visual cortex (de Pesters et al., 2016; Jensen et al., 2014; Klimesch, 2012; Stormer et al., 2016).

Of note, the weighted peak frequency of the alpha-beta suppression slightly, but significantly, increased with increasing velocity of visual motion, most probably because of stronger beta suppression associated with higher velocity (Fig. 8B). It would be interesting to investigate in more detail how activity in the beta and low gamma ranges is affected by increasing velocity of the visual motion, but this topic is beyond the scope of the present study.

### Gamma suppression transition point and E-I balance in the visual cortex

Overall, our findings indicate that the strength of excitatory drive, but not velocity tuning of visual neurons, is the main factor contributing to an initially facilitative and then suppressive effect of visual motion velocity on the gamma response power. This fact is fully consistent with predictions of computational modeling studies (Borgers and Kopell, 2005; Borgers and Walker, 2013; Cannon et al., 2014; King et al., 2013; Kopell et al., 2010), and, taken one step further, favors the idea that the gamma suppression transition point could serve as a non-invasive biomarker of the efficiency of the E/I balance regulation in the visual cortex.

It is commonly accepted that the active states of the brain related to the processing of sensory information are characterized by a ‘desynchronized’ pattern of electromagnetic activity, i.e. by a relative predominance of high-frequency (gamma) oscillations and relative suppression of those of lower frequencies (alpha, beta). Indeed, we have found that a moderate rise in sensory input drive produced an increase of gamma power, most probably due to a progressively greater recruitment of visual neurons into gamma synchronization. At first glance, the suppression of the gamma response at a yet higher level of input drive may appear counterintuitive. It has been, however predicted that gamma may desynchronize at a high level of excitatory drive due to an overexcited state of inhibitory neurons (Borgers and Kopell, 2005; King et al., 2013). It has been argued that the *asynchronous* activity of the highly excited inhibitory neurons is particularly effective in down-regulating activity in the excitatory principle cells, and therefore has an important role in homeostatic regulation of the neural E/I balance (Borgers and Kopell, 2005). In case of strong excitability of the principal cells, stronger inhibitory influences would be required to achieve a breakdown of gamma synchrony. This means that the constitutive factors that increase or decrease the excitability of the E- or I-cells may result in shifting the gamma suppression point to higher or lower intensities of excitatory drive.

Specifically, the transition to suppression of the gamma response at a relatively higher level of excitatory drive and/or the shallower slope of this suppression would reflect less effective inhibition. In indirect support of this assumption, we have recently described a lack of velocity-related gamma suppression in a subject with epilepsy and occipital spikes (Orekhova et al., 2018b), who might have an elevated E-I ratio in the visual cortex. The causal link between the E-I ratio and the gamma suppression transition point could, in the future, be proved directly using pharmacological or other (e.g. tDCS, TMS) approaches to manipulating the E-I balance.

Gamma synchronization facilitates neural communication and increases the effective output of presynaptic (Bastos et al., 2015; Fries, 2005, 2009, 2015; Ni et al., 2016; Rigoulot et al., 2017; Vinck et al., 2013). Therefore, synchronization of V1 neural activity in the gamma range is thought to enhance the effective output from the primary visual cortex to upstream cortical areas, thus affecting visual perception. A weak suppression of gamma oscillations at high stimulation intensities may then result in a stronger impact of the sensory stimulation on the neural network, and lead to sensory overload. Indeed, in a recent study, we have found that a weaker suppression of gamma response at fast motion velocities (of high-contrast gratings) correlated with enhanced sensory sensitivity, especially in the visual domain (Orekhova et al., 2018a).

Future studies in clinical populations characterized by elevated cortical excitability (e.g. epilepsy, migraine, some forms of ASD) would help to assess the potential value of gamma suppression as a biomarker for impaired regulation of the E/I balance.

## CONCLUSIONS

To summarize, our current findings provide strong support for the hypothesis linking magnitude of the visual gamma response caused by drifting gratings to the strength of excitatory drive. At the same time, our results indicate that ‘velocity turning’ of V1 neurons does not play a primary role in regulating the magnitude of the gamma response.

In both low- and high-contrast conditions the power of visual gamma oscillations demonstrated bell-shaped dependency on velocity of visual motion: it first increased with increasing velocity and then decreases with further increase in the drifting rate. As predicted by the ‘excitatory drive model’, reducing intensity of the visual input by lowering contrast of the grating shifted the maximum of the bell-shaped curve — the ‘suppression transition velocity’’ — to higher velocity values. Unlike induced gamma oscillations, the visual alpha response was not sensitive to the changes in the gratings’ velocity or contrast.

Our present findings, which support the causative role of strong excitatory drive in visual gamma response suppression, have important theoretical implications. The computational modeling studies and our previous experimental results imply that such causal relationships may reflect the capacity of inhibitory neurons to down-regulate growing excitation and to maintain the E-I balance. Therefore, the shape of the gamma response modulation curve and, in particular, the ‘suppression transition velocity’ parameter evaluated in the present study, may provide important information about regulation of the E-I balance.

We anticipate that such a combinatory index as ‘suppression transition velocity’ will be a better measure of the E-I dysfunction in brain disorders than visual gamma parameters that are extracted in a single experimental condition. Apart from having theoretical relevance, this index evades several caveats of using peak gamma amplitude and frequency. Firstly, it is inherently relational, and is therefore less sensitive to inter-individual differences in cortical anatomy and SNR. Secondly it allows to avoid ambiguity related to single-condition assessments of gamma amplitudes, which do not always correlate with each other (e.g., when measured with static vs. ‘fast movement’ conditions); Thirdly, it’s individual values have remarkable rank-order consistency when measured at different contrasts, and therefore should have high test–retest stability. Altogether, these considerations suggest that the ‘gamma suppression transition velocity’ has a translational potential as an index of the E-I balance for informing clinical practice and trials.

## Acknowledgments

This work was supported by the Moscow State University of Psychology and Education; the Charity Foundation "Way Out"; Swedish Childhood Cancer (# MT2014-0007); Knut and Alice Wallenberg Foundation (#2014.0102) and Swedish Research Council (# 2017-0068). We are grateful to all volunteer participants.

## Referenses

Alitto, H.J., Usrey, W.M., 2004. Influence of contrast on orientation and temporal frequency tuning in ferret primary visual cortex. Journal of Neurophysiology 91, 2797–2808.

Anver, H., Ward, P.D., Magony, A., Vreugdenhil, M., 2011. NMDA Receptor Hypofunction Phase Couples Independent gamma-Oscillations in the Rat Visual Cortex. Neuropsychopharmacology 36, 519–528.

Bartolo, M.J., Gieselmann, M.A., Vuksanovic, V., Hunter, D., Sun, L., Chen, X., Delicato, L.S., Thiele, A., 2011. Stimulus-induced dissociation of neuronal firing rates and local field potential gamma power and its relationship to the resonance blood oxygen level-dependent signal in macaque primary visual cortex. European Journal of Neuroscience 34, 1857–1870.

Bastos, A.M., Vezoli, J., Bosman, C.A., Schoffelen, J.M., Oostenveld, R., Dowdall, J.R., De Weerd, P., Kennedy, H., Fries, P., 2015. Visual Areas Exert Feedforward and Feedback Influences through Distinct Frequency Channels. Neuron 85, 390–401.

Borgers, C., Kopell, N., 2005. Effects of noisy drive on rhythms in networks of excitatory and inhibitory neurons. Neural Computation 17, 557–608.

Borgers, C., Walker, B., 2013. Toggling between gamma-frequency activity and suppression of cell assemblies. Front Comput Neurosc.

Bosman, C.A., Schoffelen, J.M., Brunet, N., Oostenveld, R., Bastos, A.M., Womelsdorf, T., Rubehn, B., Stieglitz, T., De Weerd, P., Fries, P., 2012. Attentional Stimulus Selection through Selective Synchronization between Monkey Visual Areas. Neuron 75, 875–888.

Buzsaki, G., Wang, X.J., 2012. Mechanisms of Gamma Oscillations. Annual Review of Neuroscience, Vol 35 35, 203–225.

Cannon, J., McCarthy, M.M., Lee, S., Lee, J., Borgers, C., Whittington, M.A., Kopell, N., 2014. Neurosystems: brain rhythms and cognitive processing. European Journal of Neuroscience 39, 705–719.

Chen, G., Zhang, Y., Li, X., Zhao, X., Ye, Q., Lin, Y., Tao, H.W., Rasch, M.J., Zhang, X., 2017. Distinct Inhibitory Circuits Orchestrate Cortical beta and gamma Band Oscillations. Neuron 96, 1403–1418 e1406.

Cousijn, H., Haegens, S., Wallis, G., Near, J., Stokes, M.G., Harrison, P.J., Nobre, A.C., 2014. Resting GABA and glutamate concentrations do not predict visual gamma frequency or amplitude. Proceedings of the National Academy of Sciences of the United States of America 111, 9301–9306.

de Pesters, A., Coon, W.G., Brunner, P., Gunduz, A., Ritaccio, A.L., Brunet, N.M., de Weerd, P., Roberts, M.J., Oostenveld, R., Fries, P., Schalk, G., 2016. Alpha power indexes task-related networks on large and small scales: A multimodal ECoG study in humans and a non-human primate. Neuroimage 134, 122–131.

Edden, R.A., Muthukumaraswamy, S.D., Freeman, T.C., Singh, K.D., 2009. Orientation discrimination performance is predicted by GABA concentration and gamma oscillation frequency in human primary visual cortex. J Neurosci 29, 15721–15726.

Ferando, I., Mody, I., 2013. Altered gamma oscillations during pregnancy through loss of delta subunit-containing GABA(A) receptors on parvalbumin interneurons. Frontiers in Neural Circuits, p. 144.

Fries, P., 2005. A mechanism for cognitive dynamics: neuronal communication through neuronal coherence. Trends in Cognitive Sciences 9, 474–480.

Fries, P., 2009. Neuronal gamma-band synchronization as a fundamental process in cortical computation. Annual Review of Neuroscience, Vol 35 32, 209–224.

Fries, P., 2015. Rhythms for Cognition: Communication through Coherence. Neuron 88, 220– 235.

Gaetz, W., Roberts, T.P.L., Singh, K.D., Muthukumaraswamy, S.D., 2012. Functional and structural correlates of the aging brain: Relating visual cortex (V1) gamma band responses to age-related structural change. Human Brain Mapping 33, 2035–2046.

Gramfort, A., Luessi, M., Larson, E., Engemann, D.A., Strohmeier, D., Brodbeck, C., Goj, R., Jas, M., Brooks, T., Parkkonen, L., Hamalainen, M., 2013. MEG and EEG data analysis with MNE-Python. Front Neurosci.

Gross, J., Kujala, J., Hamalainen, M., Timmermann, L., Schnitzler, A., Salmelin, R., 2001. Dynamic imaging of coherent sources: Studying neural interactions in the human brain. Proceedings of the National Academy of Sciences of the United States of America 98, 694–699.

Hadjipapas, A., Lowet, E., Roberts, M.J., Peter, A., De Weerd, P., 2015. Parametric variation of gamma frequency and power with luminance contrast: A comparative study of human MEG and monkey LFP and spike responses. Neuroimage 112, 327–340.

Henrie, J.A., Shapley, R., 2005. LFP power spectra in V1 cortex: the graded effect of stimulus contrast. J Neurophysiol 94, 479–490.

Hoogenboom, N., Schoffelen, J.M., Oostenveld, R., Parkes, L.M., Fries, P., 2006. Localizing human visual gamma-band activity in frequency, time and space. Neuroimage 29, 764–773.

Jensen, O., Gips, B., Bergmann, T.O., Bonnefond, M., 2014. Temporal coding organized by coupled alpha and gamma oscillations prioritize visual processing. Trends Neurosci 37, 357–369.

Jia, X.X., Smith, M.A., Kohn, A., 2011. Stimulus Selectivity and Spatial Coherence of Gamma Components of the Local Field Potential. J Neurosci 31, 9390–9403.

Jia, X.X., Xing, D.J., Kohn, A., 2013. No Consistent Relationship between Gamma Power and Peak Frequency in Macaque Primary Visual Cortex. J Neurosci 33, 17–U421.

King, P.D., Zylberberg, J., DeWeese, M.R., 2013. Inhibitory interneurons decorrelate excitatory cells to drive sparse code formation in a spiking model of V1. J Neurosci 33, 5475–5485.

Klimesch, W., 2012. alpha-band oscillations, attention, and controlled access to stored information. Trends Cogn Sci 16, 606–617.

Koelewijn, L., Dumont, J.R., Muthukumaraswamy, S.D., Rich, A.N., Singh, K.D., 2011. Induced and evoked neural correlates of orientation selectivity in human visual cortex. Neuroimage 54, 2983–2993.

Kopell, N., Börgers, C., Pervouchine, D., Malerba, P., Tort, A., 2010. Gamma and Theta Rhythms in Biophysical Models of Hippocampal Circuits. Springer Series in Computational Neuroscience Springer Science+Business Media.

Kujala, J., Jung, J., Bouvard, S., Lecaignard, F., Lothe, A., Bouet, R., Ciumas, C., Ryvlin, P., Jerbi, K., 2015. Gamma oscillations in V1 are correlated with GABA(A) receptor density: A multi-modal MEG and Flumazenil-PET study. Sci Rep 5, 16347.

Liu, Y., Bengson, J., Huang, H., Mangun, G.R., Ding, M., 2016. Top-down Modulation of Neural Activity in Anticipatory Visual Attention: Control Mechanisms Revealed by Simultaneous EEG-fMRI. Cereb Cortex 26, 517–529.

Livingstone, M.S., Conway, B.R., 2007. Contrast affects speed tuning, space-time slant, and receptive-field organization of simple cells in macaque V1. J Neurophysiol 97, 849–857.

Lowet, E., Roberts, M., Hadjipapas, A., Peter, A., van der Eerden, J., De Weerd, P., 2015. Input-Dependent Frequency Modulation of Cortical Gamma Oscillations Shapes Spatial Synchronization and Enables Phase Coding. Plos Computational Biology 11.

Mann, E.O., Mody, I., 2010. Control of hippocampal gamma oscillation frequency by tonic inhibition and excitation of interneurons. Nature Neuroscience 13, 205–U290.

Murty, D., Shirhatti, V., Ravishankar, P., Ray, S., 2018. Large Visual Stimuli Induce Two Distinct Gamma Oscillations in Primate Visual Cortex. J Neurosci 38, 2730–2744.

Muthukumaraswamy, S.D., Routley, B., Droog, W., Singh, K.D., Hamandi, K., 2016. The effects of AMPA blockade on the spectral profile of human early visual cortex recordings studied with non-invasive MEG. Cortex 81, 266–275.

Muthukumaraswamy, S.D., Singh, K.D., 2013. Visual gamma oscillations: The effects of stimulus type, visual field coverage and stimulus motion on MEG and EEG recordings. Neuroimage 69, 223–230.

Muthukumaraswamy, S.D., Singh, K.D., Swettenham, J.B., Jones, D.K., 2010. Visual gamma oscillations and evoked responses: Variability, repeatability and structural MRI correlates. Neuroimage 49, 3349–3357.

Ni, J.G., Wunderle, T., Lewis, C.M., Desimone, R., Diester, I., Fries, P., 2016. Gamma-Rhythmic Gain Modulation. Neuron 92, 240–251.

Nichols, T.E., Holmes, A.P., 2002. Nonparametric permutation tests for functional neuroimaging: A primer with examples. Human Brain Mapping 15, 1–25.

Oostenveld, R., Fries, P., Maris, E., Schoffelen, J.M., 2011. FieldTrip: Open source software for advanced analysis of MEG, EEG, and invasive electrophysiological data. Comput Intell Neurosci.

Orekhova, E.V., Butorina, A.V., Sysoeva, O.V., Prokofyev, A.O., Nikolaeva, A.Y., Stroganova, T.A., 2015. Frequency of gamma oscillations in humans is modulated by velocity of visual motion. Journal of Neurophysiology 114, 244–255.

Orekhova, E.V., Stroganova, T.A., Schneiderman, J.F., Lundström, S., Riaz, B., Sarovoc, D., Sysoeva, O.V., Brant, G., Gillberg, C., Hadjikhani, N., 2018a. Neural gain control measured through cortical gamma oscillations is associated with sensory sensitivity. Hum Brain Mapp, in press.

Orekhova, E.V., Sysoeva, O.V., Schneiderman, J.F., Lundstrom, S., Galuta, I.A., Goiaeva, D.E., Prokofyev, A.O., Riaz, B., Keeler, C., Hadjikhani, N., Gillberg, C., Stroganova, T.A., 2018b. Input-dependent modulation of MEG gamma oscillations reflects gain control in the visual cortex. Sci Rep 8, 8451.

Palva, S., Palva, J.M., 2011. Functional roles of alpha-band phase synchronization in local and large-scale cortical networks. Front Psychol 2, 204.

Perry, G., Randle, J.M., Koelewijn, L., Routley, B.C., Singh, K.D., 2015. Linear Tuning of Gamma Amplitude and Frequency to Luminance Contrast: Evidence from a Continuous Mapping Paradigm. PLoS One.

Pfurtscheller, G., da Silva, F.H.L., 1999. Event-related EEG/MEG synchronization and desynchronization: basic principles. Clinical Neurophysiology 110, 1842–1857.

Priebe, N.J., Lisberger, S.G., Movshon, J.A., 2006. Tuning for spatiotemporal frequency and speed in directionally selective neurons of macaque striate cortex. Journal of Neuroscience 26, 2941–2950.

Ray, S., Maunsell, J.H., 2010. Differences in gamma frequencies across visual cortex restrict their possible use in computation. Neuron 67, 885–896.

Reynolds, J.H., Chelazzi, L., 2004. Attentional modulation of visual processing. Annual Review of Neuroscience 27, 611–647.

Rigoulot, S., Knoth, I.S., Lafontaine, M.P., Vannasing, P., Major, P., Jacquemont, S., Michaud, J.L., Jerbi, K., Lippe, S., 2017. Altered visual repetition suppression in Fragile X Syndrome: New evidence from ERPs and oscillatory activity. Int J Dev Neurosci 59, 52–59.

Roberts, M.J., Lowet, E., Brunet, N.M., Ter Wal, M., Tiesinga, P., Fries, P., De Weerd, P., 2013. Robust Gamma Coherence between Macaque V1 and V2 by Dynamic Frequency Matching. Neuron 78, 523–536.

Romei, V., Brodbeck, V., Michel, C., Amedi, A., Pascual-Leone, A., Thut, G., 2008. Spontaneous fluctuations in posterior alpha-band EEG activity reflect variability in excitability of human visual areas. Cereb Cortex 18, 2010–2018.

Salelkar, S., Somasekhar, G.M., Ray, S., 2018. Distinct frequency bands in the local field potential are differently tuned to stimulus drift rate. J Neurophysiol.

Scheeringa, R., Koopmans, P.J., van Mourik, T., Jensen, O., Norris, D.G., 2016. The relationship between oscillatory EEG activity and the laminar-specific BOLD signal. Proceedings of the National Academy of Sciences of the United States of America 113, 6761–6766.

Schwarzkopf, D.S., Robertson, D.J., Song, C., Barnes, G.R., Rees, G., 2012. The frequency of visually induced gamma-band oscillations depends on the size of early human visual cortex. J Neurosci 32, 1507–1512.

Shapley, R.M., Victor, J.D., 1978. The effect of contrast on the transfer properties of cat retinal ganglion cells. J Physiol 285, 275–298.

Shapley, R.M., Victor, J.D., 1981. How the contrast gain control modifies the frequency responses of cat retinal ganglion cells. J Physiol 318, 161–179.

Snyder, A.C., Foxe, J.J., 2010. Anticipatory attentional suppression of visual features indexed by oscillatory alpha-band power increases: a high-density electrical mapping study. J Neurosci 30, 4024–4032.

Stormer, V., Feng, W., Martinez, A., McDonald, J., Hillyard, S., 2016. Salient, Irrelevant Sounds Reflexively Induce Alpha Rhythm Desynchronization in Parallel with Slow Potential Shifts in Visual Cortex. J Cogn Neurosci 28, 433–445.

Sumner, R.L., McMillan, R.L., Shaw, A.D., Singh, K.D., Sundram, F., Muthukumaraswamy, S.D., 2018. Peak visual gamma frequency is modified across the healthy menstrual cycle. Human Brain Mapping 39, 3187–3202.

Swettenham, J.B., Muthukumaraswamy, S.D., Singh, K.D., 2009. Spectral Properties of Induced and Evoked Gamma Oscillations in Human Early Visual Cortex to Moving and Stationary Stimuli. Journal of Neurophysiology 102, 1241–1253.

Tan, H.R.M., Gross, J., Uhlhaas, P.J., 2016. MEG sensor and source measures of visually induced gamma-band oscillations are highly reliable. Neuroimage 137, 34–44.

Taulu, S., Hari, R., 2009. Removal of magnetoencephalographic artifacts with temporal signal-space separation: demonstration with single-trial auditory-evoked responses. Hum Brain Mapp 30, 1524–1534.

Tzourio-Mazoyer, N., Landeau, B., Papathanassiou, D., Crivello, F., Etard, O., Delcroix, N., Mazoyer, B., Joliot, M., 2002. Automated anatomical labeling of activations in SPM using a macroscopic anatomical parcellation of the MNI MRI single-subject brain. Neuroimage 15, 273–289.

Van Hooser, S.D., Roy, A., Rhodes, H.J., Culp, J.H., Fitzpatrick, D., 2013. Transformation of Receptive Field Properties from Lateral Geniculate Nucleus to Superficial V1 in the Tree Shrew. Journal of Neuroscience 33, 11494–11505.

van Kerkoerle, T., Self, M.W., Dagnino, B., Gariel-Mathis, M.A., Poort, J., van der Togt, C., Roelfsema, P.R., 2014. Alpha and gamma oscillations characterize feedback and feedforward processing in monkey visual cortex. Proc Natl Acad Sci U S A 111, 14332– 14341.

van Pelt, S., Boomsma, D.I., Fries, P., 2012. Magnetoencephalography in Twins Reveals a Strong Genetic Determination of the Peak Frequency of Visually Induced Gamma-Band Synchronization. Journal of Neuroscience 32, 3388–3392.

van Pelt, S., Shumskaya, E., Fries, P., 2018. Cortical volume and sex influence visual gamma. Neuroimage 178, 702–712.

VanVeen, B.D., vanDrongelen, W., Yuchtman, M., Suzuki, A., 1997. Localization of brain electrical activity via linearly constrained minimum variance spatial filtering. Ieee Transactions on Biomedical Engineering 44, 867–880.

Vinck, M., Womelsdorf, T., Fries, P., 2013. Gamma-band synchronization and information transmission. In: Quiroga, R.Q., Panzeri, S. (Eds.), Principles of Neural Coding. CRC Press, pp. 449–469.

Yuan, H., Liu, T., Szarkowski, R., Rios, C., Ashe, J., He, B., 2010. Negative covariation between task-related responses in alpha/beta-band activity and BOLD in human sensorimotor cortex: an EEG and fMRI study of motor imagery and movements. Neuroimage 49, 2596–2606.

