## Supplementary Table for "Additive effect of contrast and velocity proves the role of strong excitatory drive in suppression of visual gamma response"

**Supplementary Tables**

**Supplementary table 1A.** The 100% contrast: position of the ‘maximally induced gamma voxel’ in the ‘static’, ‘slow’, ‘medium’ and ‘fast’ velocity conditions in each of the 17 participants. The results are presented only in case of significant brain cluster of gamma increase, as well as significant increase of the peak gamma power in the 25 voxels selection (p<0.0001).

|  | | Area of the ‘maximally induced voxel’** | | | | MPI coordinates of the ‘maximally induced voxel’ (cm) | | | | | | | | | | | | | | | |
| --- | --- | --- | --- | --- | --- | --- | --- | --- | --- | --- | --- | --- | --- | --- | --- | --- | --- | --- | --- | --- | --- |
| Subj | Static | | Slow | Medium | Fast | Static | | | | Slow | | | | Medium | | | | Fast | | | |
|  |  |  |  |  |  | | X0 | Y0 | Z0 | | X 1 | Y1 | Z1 | | X2 | Y2 | Z2 | | X3 | Y3 | Z3 |
| 1 | Calcarine_L | | Calcarine_L | Calcarine_L | Calcarine_L | | 0.2 | -10 | 0.2 | | 0.2 | -10 | 0.8 | | 0.2 | -10 | 0.8 | | 0.2 | -10 | 0.8 |
| 2 | Calcarine_L | | Calcarine_R | Calcarine_R | Calcarine_L | | 0.2 | -8.8 | 0.8 | | 0.8 | -8.8 | 0.2 | | 0.8 | -8.8 | 0.2 | | 0.2 | -8.8 | 0.8 |
| 3 | Calcarine_L | | Occip_Sup_L | Calcarine_L | Calcarine_L | | -0.4 | -8.8 | -1 | | -1 | -9.4 | 0.8 | | -0.4 | -8.8 | 0.8 | | -0.4 | -8.8 | 0.8 |
| 4 | Cuneus_R | | Cuneus_R | Cuneus_R | Cuneus_L | | 2 | -9.4 | 2 | | 2 | -9.4 | 0.8 | | 2 | -9.4 | 0.8 | | -0.4 | -9.4 | 1.4 |
| 5 | Cuneus_R | | Cuneus_R | Cuneus_R | Cuneus_R | | 1.4 | -10 | 1.4 | | 1.4 | -10 | 0.8 | | 1.4 | -10 | 0.8 | | 1.4 | -9.4 | 0.8 |
| 6 | Calcarine_L | | Calcarine_L | Calcarine_L | Calcarine_L | | -0.4 | -10 | -0.4 | | -0.4 | -10 | 0.2 | | -0.4 | -9.4 | 0.2 | | 0.2 | -8.8 | 0.8 |
| 7 | Calcarine_R | | Calcarine_R | Calcarine_R | Calcarine_R | | 1.4 | -8.8 | 0.8 | | 0.8 | -8.8 | 0.8 | | 0.8 | -8.8 | 0.8 | | 0.8 | -8.8 | 0.8 |
| 8 | Cuneus_L | | Occip_Sup_R | Cuneus_R | Occip_Sup R | | -1 | -9.4 | 2.6 | | 2.6 | -9.4 | 1.4 | | 2 | -9.4 | 1.4 | | 2 | -9.4 | 2 |
| 9 | Calcarine_L | | Calcarine_L | Calcarine_L | Calcarine_L | | 0.2 | -9.4 | 0.2 | | 0.2 | -9.4 | -0.4 | | 0.2 | -9.4 | 0.2 | | 0.2 | -9.4 | -0.4 |
| 10 | Occip_Mid_L | | Occip_Mid_L | Occip_Mid_L | Occip_Mid_L | | -1.6 | -9.4 | -1.6 | | -1.6 | -9.4 | 0.2 | | -1.6 | -9.4 | 0.2 | | -2.2 | -10 | 0.8 |
| 11 | Calcarine_L | | Calcarine_R | Lingual_R | Lingual_L | | -0.4 | -8.8 | 0.8 | | 0.8 | -8.8 | 0.2 | | 0.8 | -8.8 | -0.4 | | -0.4 | -8.2 | 0.2 |
| 12 | Calcarine_L | | Calcarine_L | Calcarine_L | Calcarine_L | | 0.2 | -10 | 0.2 | | 0.2 | -10 | -0.4 | | 0.2 | -9.4 | -0.4 | | -0.4 | -9.4 | 0.2 |
| 13 | Occip_Sup_L | | Cuneus_R | Occip_Sup_L | Occip_Sup_L | | -1.6 | -9.4 | 1.4 | | 1.4 | -10 | 0.8 | | -1 | -9.4 | 0.8 | | -1.6 | -9.4 | 1.4 |
| 14 | Calcarine_L | | Calcarine_R | Calcarine_R | Calcarine_R | | -0.4 | -9.4 | 1.4 | | 1.4 | -8.8 | 1.4 | | 0.8 | -8.8 | 0.8 | | 1.4 | -8.8 | 1.4 |
| 15 | Cuneus_L | | Cuneus_L | Cuneus_L | Cuneus_L | | -0.4 | -9.4 | -0.4 | | -0.4 | -9.4 | 2 | | 0.2 | -9.4 | 2 | | 0.2 | -9.4 | 1.4 |
| 16 | Calcarine_L* | | Calcarine_L | Calcarine_L | Calcarine_L | | -0.4 | -9.4 | 0.2 | | 0.2 | -9.4 | -1 | | -0.4 | -9.4 | -0.4 | | -0.4 | -9.4 | -0.4 |
| 17 | Occip_Mid_L | | Occip_Mid_L | Occip_Mid_L | Calcarine_L | | -1 | -10 | -1 | | -1 | -10 | 0.2 | | -1 | -10 | 0.2 | | -0.4 | -9.4 | 0.8 |

* According to LCMV analysis the maximal gamma increase occurred outside visual cortical areas. The DICS beamformer analysis, however, revealed a significant cluster of gamma increase locally in the visual cortex. In this case the maximally induced voxel was still chosen within visual cortical areas.

** According to AAL atlas (N. Tzourio-Mazoyer, B. Landeau, D. Papathanassiou, F. Crivello, O. Etard, N. Delcroix, B. Mazoyer, and M. Joliot. Automated Anatomical Labeling of Activations in SPM Using a Macroscopic Anatomical Parcellation of the MNI MRI Single-Subject Brain. NeuroImage 2002. 15:273-289.)

**Supplementary table 1B.** The 50% contrast: position of the ‘maximally induced gamma voxel’ in the ‘static’. ‘slow’. ‘medium’ and ‘fast’ velocity conditions in each of the 17 participants. The results are presented only in case of significant brain cluster of gamma increase, as well as significant increase of the peak gamma power in the 25 voxels selection (p<0.0001).

|  | | Location of the occipital ‘maximally induced voxel’ | | | | MPI coordinates of the ‘maximally induced voxel’ | | | | | | | | | | | | | | | |
| --- | --- | --- | --- | --- | --- | --- | --- | --- | --- | --- | --- | --- | --- | --- | --- | --- | --- | --- | --- | --- | --- |
| Subj | Static | | Slow | Medium | Fast | Static | | | | Slow | | | | Medium | | | | Fast | | | |
|  |  |  |  |  |  | | X0 | Y0 | Z0 | | X 1 | Y1 | Z1 | | X2 | Y2 | Z2 | | X3 | Y3 | Z3 |
| 1 | Calcarine_L | | Calcarine_L | Calcarine_L | Calcarine_L | | 0.2 | -10 | 0.8 | | 0.2 | -10 | 0.8 | | 0.2 | -10 | 0.8 | | 0.2 | -10 | 0.8 |
| 2 | Calcarine_L | | Calcarine_L | Calcarine_L | Calcarine_L | | -0.4 | -9.4 | 0.2 | | 0.2 | -8.8 | 0.2 | | -0.4 | -8.8 | 0.8 | | -0.4 | -8.8 | 0.8 |
| 3 | Cuneus_R* | | Calcarine_L | Calcarine_L | Calcarine_L | | 2 | -8.8 | 1.4 | | -0.4 | -8.8 | 0.8 | | -0.4 | -8.8 | 0.2 | | -0.4 | -8.8 | 0.2 |
| 4 | Cuneus_L | | Cuneus_R | Cuneus_R | Cuneus_L | | 0.2 | -9.4 | 1.4 | | 2 | -9.4 | 1.4 | | 2 | -9.4 | 0.8 | | 0.2 | -9.4 | 2 |
| 5 | Cuneus_R | | Cuneus_R | Cuneus_R | Cuneus_R | | 1.4 | -9.4 | 0.8 | | 1.4 | -10 | 0.8 | | 1.4 | -9.4 | 0.8 | | 1.4 | -10 | 1.4 |
| 6 | Calcarine_L | | Calcarine_L | Calcarine_L | Calcarine_R | | -0.4 | -10 | 0.8 | | -0.4 | -10 | 0.8 | | -0.4 | -9.4 | 0.2 | | 0.8 | -9.4 | 0.8 |
| 7 |  | |  |  |  | |  |  |  | |  |  |  | |  |  |  | |  |  |  |
| 8 |  | | Cuneus_R | Occip_Sup_R |  | |  |  |  | | 2 | -9.4 | 1.4 | | 2.6 | -9.4 | 1.4 | |  |  |  |
| 9 | Calcarine_L | | Calcarine_L | Calcarine_L | Calcarine_L | | 0.2 | -9.4 | -0.4 | | 0.2 | -9.4 | -0.4 | | 0.2 | -9.4 | 0.2 | | 0.2 | -9.4 | 0.2 |
| 10 | Calcarine_R | | Occil_Mid_L | Occip_Mid_L | Occip_Mid_L | | 0.8 | -9.4 | 0.8 | | -1.6 | -9.4 | 0.2 | | -1.6 | -9.4 | 0.2 | | -2.2 | -10 | 0.8 |
| 11 | Lingual_L | | Lingual_R | Calcarine_L | Calcarine_R | | -1 | -8.2 | -0.4 | | 0.8 | -8.8 | -0.4 | | -1 | -8.8 | 0.2 | | 1.4 | -9.4 | -0.4 |
| 12 | Occip_Mid_L | | Calcarine_L | Calcarine_L | Occip_Sup_L | | -1 | -10 | 0.2 | | -0.4 | -9.4 | 0.2 | | -0.4 | -9.4 | 0.2 | | -1 | -9.4 | 0.2 |
| 13 | Cuneus_L | | Occip_Sup_L | Cuneus_L | Cuneus_L | | 0.2 | -10 | 1.4 | | -1.6 | -10 | 1.4 | | 0.2 | -10 | 1.4 | | 0.2 | -10 | 1.4 |
| 14 | Calcarine_L | | Calcarine_L | Lingual_R | Calcarine_L | | -0.4 | -10 | 0.2 | | -0.4 | -9.4 | -0.4 | | 0.8 | -8.8 | -0.4 | | -0.4 | -9.4 | 0.2 |
| 15 | Cuneus_L | | Cuneus_L | Cuneus_L | Occip_Sup_L | | -0.4 | -9.4 | 2 | | -0.4 | -9.4 | 2 | | -0.4 | -9.4 | 2 | | -0.4 | -10 | 2 |
| 16 |  | | Calcarine_L | Calcarine_L | Calcarine_L | |  |  |  | | -0.4 | -9.4 | -0.4 | | -1 | -9.4 | -1 | | 0.2 | -9.4 | -1 |
| 17 | Occip_Sup_L | | Occip_Mid_L | Occip_Mid_L | Cuneus_L | | -1 | -10 | 1.4 | | -1 | -10 | 0.2 | | -1 | -10 | 0.2 | | -0.4 | -9.4 | 1.4 |

* According to LCMV analysis the maximal gamma increase occurred outside visual cortical areas. The DICS beamformer analysis, however, revealed a significant cluster of gamma increase locally in the visual cortex. In this case the maximally induced voxel was still chosen within visual cortical areas.

**Supplementary table 2A.** The 100% contrast: position of the ‘maximal alpha-beta suppression voxel’ in the ‘static’, ‘slow’, ‘medium’ and ‘fast’ velocity conditions in each of the 17 participants. The results are presented only in case of significant brain cluster of alpha-beta suppression.

|  | | Area of the ‘maximally induced voxel’ | | | | MPI coordinates of the ‘maximally induced voxel’ (cm) | | | | | | | | | | | | | | | |
| --- | --- | --- | --- | --- | --- | --- | --- | --- | --- | --- | --- | --- | --- | --- | --- | --- | --- | --- | --- | --- | --- |
| Subj | Static | | Slow | Medium | Fast | Static | | | | Slow | | | | Medium | | | | Fast | | | |
|  |  |  |  |  |  | | X0 | Y0 | Z0 | | X 1 | Y1 | Z1 | | X2 | Y2 | Z2 | | X3 | Y3 | Z3 |
| 1 | Pariet_Sup_L | | Pariet_Sup_L | Pariet_Sup_L | Occip_Mid_L | | -2.8 | -9.4 | -0.4 | | -1 | -10.6 | 0.2 | | -1.6 | -10 | 0.8 | | -1 | -10.6 | 0.2 |
| 2 |  | | Occip_Mid_R | Cuneus_R | Calcarine_R | |  |  |  | | 2.6 | -9.4 | 0.8 | | 2 | -8.8 | 0.8 | | 2.6 | -9.4 | 0.2 |
| 3 | Occip_Mid_R | | Occip_Inf_R | Occip_Inf_L | Occip_Mid_L | | 3.2 | -9.4 | 0.2 | | 4.4 | -8.2 | -1 | | -3.4 | -8.8 | -1 | | -3.4 | -8.8 | -0.4 |
| 4 | Occip_Mid_R | | Calcarine_L | Occip_Sup_R | Occip_Inf_R | | 3.2 | -9.4 | 1.4 | | -1 | -10 | -0.4 | | 2.6 | -9.4 | 1.4 | | 3.8 | -9.4 | -0.4 |
| 5 | Calcarine_L | | Lingual_L | Calcarine_L | Calcarine_L | | 0.2 | -9.4 | 0.8 | | -1.6 | -9.4 | -1.6 | | -0.4 | -10 | 0.8 | | -0.4 | -10 | 0.8 |
| 6 | Cuneus_R | | Occip_Sup_R | Occip_Sup_L | Cuneus_L | | 1.4 | -8.2 | 2 | | 2 | -8.2 | 2 | | -1.6 | -9.4 | 1.4 | | -1 | -9.4 | 1.4 |
| 7 | Cerebelum R | | Cerebelum R | Occip_Inf_L | Cerebelum R | | -1.6 | -10 | -1.6 | | -1.6 | -10 | -1.6 | | -1.6 | -9.4 | -1 | | 2.6 | -9.4 | 0.8 |
| 8 | Occip_Mid_R | | Occip_Mid_L | Occip_Mid_R | Occip_Sup_L | | 4.4 | -8.2 | 2 | | -2.8 | -9.4 | 2 | | 4.4 | -8.2 | 2 | | -2.8 | -9.4 | 2.6 |
| 9 | Occip_Mid_R | | Occip_Inf_R | Occip_Inf_R | Occip_Inf_L | | 3.8 | -8.8 | 0.2 | | 3.2 | -8.2 | -1.6 | | 3.2 | -8.8 | -1.6 | | -2.2 | -9.4 | -0.4 |
| 10 | Occip_Mid_R | | Calcarine_L | Occip_Mid_R | Occip_Mid_R | | 2.6 | -9.4 | 0.8 | | 0.2 | -10 | 0.2 | | 3.8 | -9.4 | 0.8 | | 2.6 | -9.4 | 0.8 |
| 11 | Calcarine_R | | Calcarine_R | Calcarine_R | Calcarine_R | | 1.4 | -9.4 | 0.2 | | 2 | -9.4 | 0.2 | | 1.4 | -9.4 | 0.2 | | 1.4 | -9.4 | 0.2 |
| 12 | Occip_Mid_L | | Calcarine_R | Occip_Mid_L | Occip_Mid_L | | -1.6 | -10 | 0.8 | | 2 | -10 | 0.2 | | -2.2 | -10 | 0.8 | | -1.6 | -10 | 0.8 |
| 13 | Occip_Mid_R | | Calcarine_L | Occip_Mid_R | Occip_Mid_L | | 4.4 | -8.2 | 2.6 | | -0.4 | -10.6 | -0.4 | | 4.4 | -8.2 | 2 | | -2.8 | -9.4 | 1.4 |
| 14 | Lingual_L | | Occip_Sup_R | Lingual_L | Occip_Mid_R | | -1.6 | -10 | -1.6 | | 2.6 | -9.4 | 1.4 | | -1.6 | -10 | -1.6 | | 3.2 | -9.4 | 1.4 |
| 15 | Temp_Mid_R | | Occip_Mid_R | Cuneus_R | Occip_Mid_R | | 4.4 | -7.6 | 0.8 | | 4.4 | -7.6 | 1.4 | | 2 | -10 | 0.8 | | 4.4 | -7.6 | 0.8 |
| 16 | Occip_Inf_L | | Lingual_L | Lingual_L | Calcarine_L | | -2.2 | -8.8 | -1 | | -1.6 | -8.8 | -1 | | -1.6 | -8.8 | -1 | | -1 | -8.8 | -0.4 |
| 17 |  | |  |  |  | |  |  |  | |  |  |  | |  |  |  | |  |  |  |

**Supplementary table 2B.** The 50% contrast: position of the ‘maximal alpha-beta suppression voxel’ in the ‘static’, ‘slow’, ‘medium’ and ‘fast’ velocity conditions in each of the 17 participants. The results are presented only in case of significant brain cluster of alpha-beta suppression.

|  | | Area of the ‘maximally induced voxel’ | | | | MPI coordinates of the ‘maximally induced voxel’ (cm) | | | | | | | | | | | | | | | |
| --- | --- | --- | --- | --- | --- | --- | --- | --- | --- | --- | --- | --- | --- | --- | --- | --- | --- | --- | --- | --- | --- |
| Subj | Static | | Slow | Medium | Fast | Static | | | | Slow | | | | Medium | | | | Fast | | | |
|  |  |  |  |  |  | | X0 | Y0 | Z0 | | X 1 | Y1 | Z1 | | X2 | Y2 | Z2 | | X3 | Y3 | Z3 |
| 1 | Pariet_Sup_L | | Pariet_Sup_L | Pariet_Sup_L | Occip_Mid_L | | -2.8 | -9.4 | -0.4 | | -1 | -10.6 | 0.2 | | -1.6 | -10 | 0.8 | | -1 | -10.6 | 0.2 |
| 2 |  | | Occip_Mid_R | Cuneus_R | Calcarine_R | |  |  |  | | 2.6 | -9.4 | 0.8 | | 2 | -8.8 | 0.8 | | 2.6 | -9.4 | 0.2 |
| 3 | Occip_Mid_R | |  | Occip_Inf_L | Occip_Mid_L | | 3.2 | -9.4 | 0.2 | |  |  |  | | -3.4 | -8.8 | -1 | | -3.4 | -8.8 | -0.4 |
| 4 | Occip_Mid_R | | Calcarine_L | Occip_Sup_R | Occip_Inf_R | | 3.2 | -9.4 | 1.4 | | -1 | -10 | -0.4 | | 2.6 | -9.4 | 1.4 | | 3.8 | -9.4 | -0.4 |
| 5 | Calcarine_L | | Lingual_L | Calcarine_L | Calcarine_L | | 0.2 | -9.4 | 0.8 | | -1.6 | -9.4 | -1.6 | | -0.4 | -10 | 0.8 | | -0.4 | -10 | 0.8 |
| 6 | Cuneus_R | | Occip_Sup_R | Occip_Sup_L | Cuneus_L | | 1.4 | -8.2 | 2 | | 2 | -8.2 | 2 | | -1.6 | -9.4 | 1.4 | | -1 | -9.4 | 1.4 |
| 7 | Cerebelum R | | Cerebelum R | Occip_Inf_L | Cerebelum R | | -1.6 | -10 | -1.6 | | -1.6 | -10 | -1.6 | | -1.6 | -9.4 | -1 | | 2.6 | -9.4 | 0.8 |
| 8 | Occip_Mid_R | | Occip_Mid_L | Occip_Mid_R | Occip_Sup_L | | 4.4 | -8.2 | 2 | | -2.8 | -9.4 | 2 | | 4.4 | -8.2 | 2 | | -2.8 | -9.4 | 2.6 |
| 9 | Occip_Mid_R | | Occip_Inf_R | Occip_Inf_R | Occip_Inf_L | | 3.8 | -8.8 | 0.2 | | 3.2 | -8.2 | -1.6 | | 3.2 | -8.8 | -1.6 | | -2.2 | -9.4 | -0.4 |
| 10 | Occip_Mid_R | | Calcarine_L | Occip_Mid_R | Occip_Mid_R | | 2.6 | -9.4 | 0.8 | | 0.2 | -10 | 0.2 | | 3.8 | -9.4 | 0.8 | | 2.6 | -9.4 | 0.8 |
| 11 | Calcarine_R | | Calcarine_R | Calcarine_R | Calcarine_R | | 1.4 | -9.4 | 0.2 | | 2 | -9.4 | 0.2 | | 1.4 | -9.4 | 0.2 | | 1.4 | -9.4 | 0.2 |
| 12 |  | | Calcarine_R | Occip_Mid_L | Occip_Mid_L | |  |  |  | | 2 | -10 | 0.2 | | -2.2 | -10 | 0.8 | | -1.6 | -10 | 0.8 |
| 13 | Occip_Mid_R | | Calcarine_L | Occip_Mid_R | Occip_Mid_L | | 4.4 | -8.2 | 2.6 | | -0.4 | -10.6 | -0.4 | | 4.4 | -8.2 | 2 | | -2.8 | -9.4 | 1.4 |
| 14 | Lingual_L | | Occip_Sup_R | Lingual_L | Occip_Mid_R | | -1.6 | -10 | -1.6 | | 2.6 | -9.4 | 1.4 | | -1.6 | -10 | -1.6 | | 3.2 | -9.4 | 1.4 |
| 15 | Temp_Mid_R | | Occip_Mid_R | Cuneus_R | Occip_Mid_R | | 4.4 | -7.6 | 0.8 | | 4.4 | -7.6 | 1.4 | | 2 | -10 | 0.8 | | 4.4 | -7.6 | 0.8 |
| 16 | Occip_Inf_L | | Lingual_L | Lingual_L | Calcarine_L | | -2.2 | -8.8 | -1 | | -1.6 | -8.8 | -1 | | -1.6 | -8.8 | -1 | | -1 | -8.8 | -0.4 |
| 17 | Occipi_Inf_R | | Calcarine_R | Occip_Sup_R | Calcarine_R | | 2 | -10 | 0.2 | | 3.8 | -9.4 | -1 | | -1.6 | -10 | 0.8 | | 2 | -10 | 0.8 |

**Supplementary table 3.** Partial correlations between peak frequencies of gamma responses elicited by visual gratings drifting with different velocities in the two contrast conditions. Age was taken as a nuisance variable. Frequency was evaluated only for reliable gamma responses (see Methods).

|  | 100% contrast | | | |
| --- | --- | --- | --- | --- |
|  | 0°/s | 1.2°/s | 3.6°/s | 6.0°/s |
| 0°/s |  | **0,83***** | **0.65*** | 0.36 |
| 1.2°/s |  |  | **0.92***** | **0.72**** |
| 3.6°/s |  |  |  | **0.87***** |
|  | 50% contrast | | | |
|  | 0°/s | 1.2°/s | 3.6°/s | 6.0°/s |
| 0°/s |  | **0.68**** | **0.6*** | **0.58*** |
| 1.2°/s |  |  | **0.93***** | **0.84***** |
| 3.6°/s |  |  |  | **0.91***** |
