## Supplementary Figure for "Additive effect of contrast and velocity proves the role of strong excitatory drive in suppression of visual gamma response"

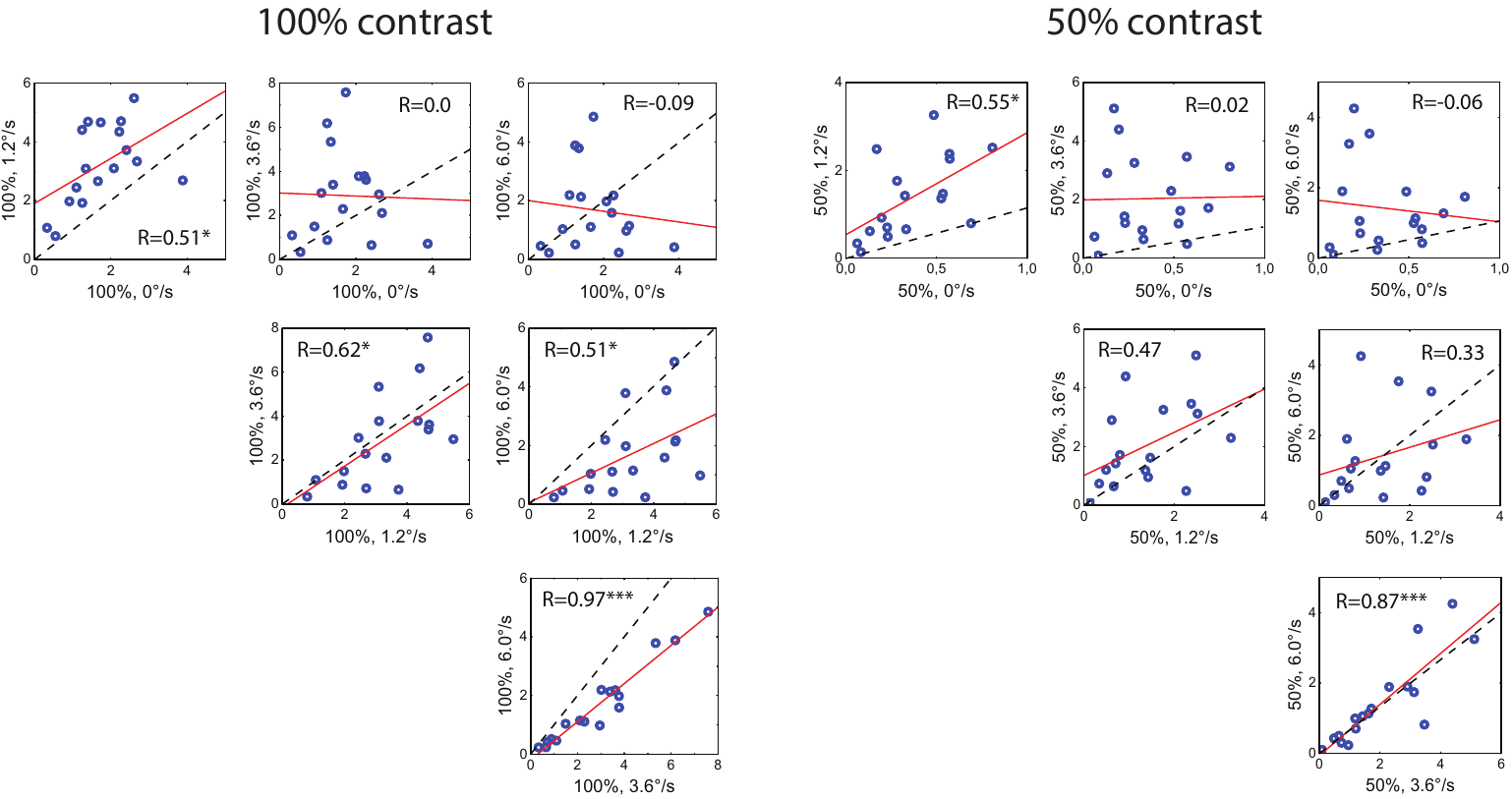

**Supplementary figure 1.** Correlations between gamma response power values measured in different velocity conditions at the 100% and 50% contrasts. Blue dots denote individual gamma response power values measured as (Pow_post_- Pow_pre_)/ Pow_pre_. The linear regression is shown in red. The dashed line corresponds to the axis of symmetry.
